# Chromosome-autonomous feedback downregulates meiotic DSB competence upon synaptonemal complex formation

**DOI:** 10.1101/2020.05.11.089367

**Authors:** Xiaojing Mu, Hajime Murakami, Neeman Mohibullah, Scott Keeney

**Author notes:** Integrated Genomics Operation, Memorial Sloan Kettering Cancer Center, New York, NY 10065.

## Abstract

The number of DNA double-strand breaks (DSBs) initiating meiotic recombination is elevated in *Saccharomyces cerevisiae* mutants that are globally defective in forming crossovers and synaptonemal complex (SC), a protein scaffold juxtaposing homologous chromosomes. These mutants thus appear to lack a negative feedback loop that inhibits DSB formation when homologs engage one another. This feedback is predicted to be chromosome autonomous, but this has not been tested. Moreover, what chromosomal process is recognized as “homolog engagement” remains unclear. To address these questions, we evaluated effects of homolog engagement defects restricted to small portions of the genome using karyotypically abnormal yeast strains with a homeologous chromosome V pair, monosomic V, or trisomy XV. We found that homolog-engagement-defective chromosomes incurred more DSBs, concomitant with prolonged retention of the DSB-promoting protein Rec114, while the rest of the genome remained unaffected. SC-deficient, crossover-proficient mutants *ecm11* and *gmc2* experienced increased DSB numbers diagnostic of homolog engagement defects. These findings support the hypothesis that SC formation provokes DSB protein dissociation, leading in turn to loss of a DSB competent state. Our findings show that DSB number is regulated in a chromosome-autonomous fashion and provide insight into how homeostatic DSB controls respond to aneuploidy during meiosis.

**bioRxiv version 2 (September 2020)** We added one bioRxiv citation in the discussion section which shows increased breaks in *gmc2* and *ecm11* mutants. This work was done independently by Keun Kim, Miki Shinohara and colleagues. The pulse gel electrophoresis result shown in Figure 1D has been re-quantified. The conclusion remains that homeologous and monosomic chromosome V generate more DSBs compared to internally controlled chromosome III. One *zip3* map included in PCA (Figure 6C) and hierarchical clustering analysis (Figure S5B) has been deleted because this particular map has not yet been published. This change does not affect the conclusion. An additional supplemental table (S5) has been added to show Spo11-oligo datasets used in this study. Figures and preprint have been reformatted.

---

Programmed DSB formation by Spo11 initiates meiotic homologous recombination (Lam and Keeney 2014; Hunter 2015). A fraction of DSBs are repaired as crossovers, contributing to physical linkages essential for proper segregation of homologs in meiosis I. Although DSBs can be deleterious, meiotic cells make them in large numbers (150–200 DSBs per cell in *S. cerevisiae*) (Pan et al. 2011). To mitigate the risk of genome instability, DSB formation is controlled quantitatively, spatially, and temporally by a network of intersecting feedback circuits (Cooper et al. 2014; Keeney et al. 2014; Murakami and Keeney 2014).

One circuit involves inhibition of DSB formation after homologous chromosomes have engaged one another. In *S. cerevisiae*, mutants lacking any of several ZMM proteins exhibit elevated DSB levels (Thacker et al. 2014; Anderson et al. 2015; Marsolier-Kergoat et al. 2018). ZMM proteins (Zip1–4, Msh4–5, Mer3, Spo16 and Pph3) promote formation of type I (interfering) crossovers and assembly of SC, a proteinaceous structure that includes the aligned axes of homologs plus the central region components that hold them together (Lynn et al. 2007; Pyatnitskaya et al. 2019). The elevated DSBs in ZMM mutants suggest that DSB control is defective when either synapsis or crossing over (or both) is impaired. A similar conclusion was reached on the basis of additional DSB formation in mutants with recombination defects from altered usage of strand exchange proteins (Lao et al. 2013). In mouse spermatocytes, chromosome segments that fail to synapse—either naturally on the nonhomologous parts of the X–Y pair or in response to a recombination-defective mutation—accumulate a higher density of foci of the strand exchange protein RAD51, a cytological marker for DSBs (Kauppi et al. 2013). A prolonged period of DSB formation has also been proposed to account for elevated DSB markers in *Caenorhabditis elegans* mutants with synapsis and/ or crossover defects (Nabeshima et al. 2004; Hayashi et al. 2010; Henzel et al. 2011).

To explain the yeast findings, we proposed that a ZMM-dependent process feeds back to inhibit DSB formation (Keeney et al. 2014; Thacker et al. 2014). The molecular identity of this process was (and remains) undefined, so we used the mechanistically neutral term “homolog engagement” to describe it. One possibility is SC formation, consistent with the behavior of asynaptic chromosome segments in mice (Kauppi et al. 2013). Indeed, earlier work showed that synapsis in mice is followed by displacement from chromosomes of the DSB-promoting axis protein HORMAD1 (Wojtasz et al. 2009). This observation led to the proposal that SC formation downregulates DSB formation by removing proteins needed for SPO11 activity (Wojtasz et al. 2009), an idea independently proposed later on the basis of unsynapsed regions in yeast retaining Spo11 accessory proteins such as Rec114 (Panizza et al. 2011; Carballo et al. 2013).

A nonexclusive alternative is that some aspect of crossover formation is the trigger for ZMM-dependent feedback (Keeney et al. 2014; Thacker et al. 2014). In *C. elegans*, prolonged chromosome binding of DSB-promoting proteins DSB-1 and DSB-2 occurs in mutants that cannot make crossover-designated recombination intermediates, even if SC is formed (Rosu et al. 2013; Stamper et al. 2013). This regulation involves the kinase CHK-2 and occurs nucleus-wide in response to a crossover defect on just a single chromosome pair (Carlton et al. 2006; Rosu et al. 2013; Stamper et al. 2013). In mouse, by contrast, CHK2 does not appear to regulate DSB number (Pacheco et al. 2015) and the increase in DSB levels caused by asynapsis appears to be restricted to unsynapsed regions, i.e., is chromosome autonomous (Kauppi et al. 2013). Whether yeast DSB control more closely resembles *C. elegans* or mouse (or neither) is not yet clear.

One limitation has been that most available data in yeast are from strains with catastrophic meiotic failure (e.g., ZMM mutants) that can simultaneously impinge on multiple feedback circuits including hyper-activation of Tel1 (ATM) and Mec1 (ATR) regulated pathways (Gray et al. 2013; Cooper et al. 2014; Keeney et al. 2014; Thacker et al. 2014). To circumvent this limitation, we used here a set of karyotypically abnormal *S. cerevisiae* strains in which defects in synapsis and/or crossing over are confined to specific parts of the genome through sequence divergence or chromosome gain or loss. This experimental setting enabled us to demonstrate that feedback through homolog engagement is chromosome autonomous and is accompanied by removal of Rec114 from chromosome pairs that have engaged one another. Additionally, analysis of SC-deficient but crossover-proficient *ecm11Δ* and *gmc2Δ* mutants showed that SC formation is essential and that crossing over without SC formation is not sufficient to support feedback control of DSB numbers.

## Results

### Karyotypically abnormal S. cerevisiae strains

The karyotype abnormalities we studied and their predicted effects on homolog engagement are cartooned in **Fig. 1A**. Homolog-engagement-proficient parts of the genome serve as internal controls to counter culture-to-culture variation in meiotic timing or efficiency, making these systems sensitive enough to detect even small differences.

**Figure 1.**
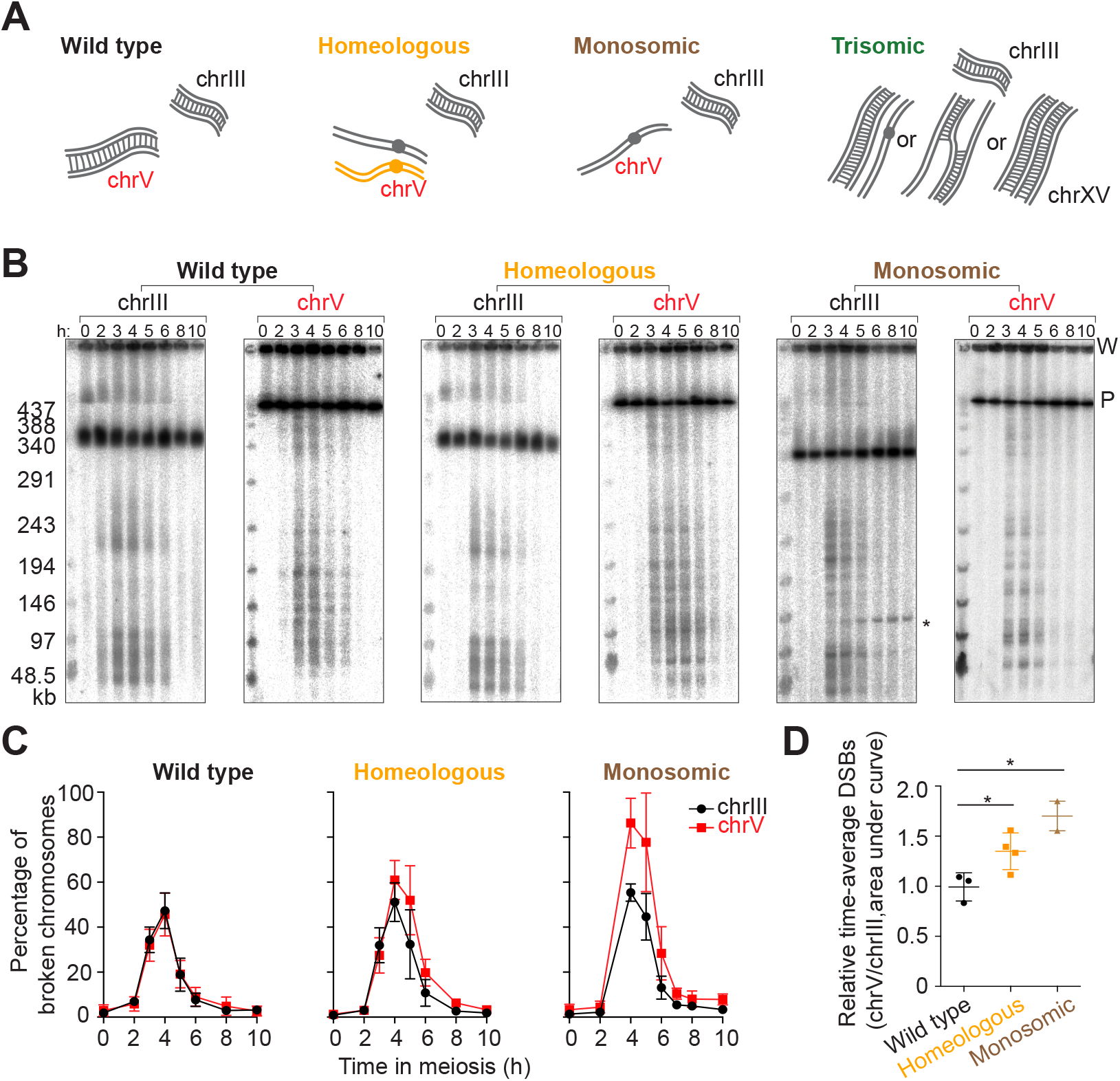
Higher DSB levels on homolog-engagement defective chromosomes. (A) Cartoons of wild-type, homeologous, monosomic and trisomic chromosome configurations. Grey lines are *S. cerevisiae* chromosomes and orange lines are *S. pastorianus*. The homeologous chrV pair rarely synapses or recombines. The trisomic chromosomes can adopt different synaptic configurations (”II+I”, partner switch, and triple synapsis). (B) Representative PFGE Southern blots probed for chrIII and chrV. P, signal from parental-length DNA; W, signal in wells. Asterisk indicates an ectopic recombination product between *leu2::hisG* (on chrIII) and *ho::hisG* (on chrIV) in the monosomic strain. The other strains do not form this product because they do not have a *hisG* insert at ho. (C) Poisson-corrected DSB quantification of PFGE Southern blots. (D) Quantification of time-averaged DSBs on chrV relative to chrIII for wild-type, homeologous and monosomic strains. * p<0.05, unpaired t test. Error bars in C, D are mean ± SD except for the monosomic strain (mean ± range).

The homeologous strain contains one copy of chromosome V (chrV) introgressed from *S. pastorianus* (also known as *S. carlsbergensis*) in an otherwise *S. cerevisiae* background (Dawson et al. 1986; Nilssontillgren et al. 1986; Goldman and Lichten 2000; Maxfield Boumil et al. 2003; Kemp et al. 2004). The homeologous chrV copies differ by about 30% from one another (Maxfield Boumil et al. 2003). As a result they show little if any evidence of homologous pairing or complete synapsis along their lengths and only very rarely produce crossovers despite making DSBs (Dawson et al. 1986; Nilssontillgren et al. 1986; Goldman and Lichten 2000; Maxfield Boumil et al. 2003; Kemp et al. 2004; Newnham et al. 2010).

The monosomic strain was generated by inducing loss of one copy of chrV before meiotic entry using a centromere-destabilizing system in which an inducible *GAL* promoter drives transcription across the centromere (Hill and Bloom 1987). We developed a presporulation procedure that yields efficient chromosome loss (**Supplemental Fig. S1** and **Supplemental Table S1**) (see Materials and Methods).

Unlike *zip3* mutants (Agarwal and Roeder 2000; Borner et al. 2004), the homeologous and monosomic strains progressed through meiosis without a strong arrest in prophase I (**Supplemental Fig. S2A**). However, both strains exhibited a moderate delay (~2 h) in nuclear division, likely from the spindle assembly checkpoint responding to non-exchange chromosomes (Shonn et al. 2000; Marston and Wassmann 2017; Lane and Kauppi 2019). The DSBs on these chromosomes are presumably repaired by recombination between sister chromatids (de Massy et al. 1994; Kearney et al. 2001; Hochwagen et al. 2005; Goldfarb and Lichten 2010).

We also present data from a serendipitously generated strain bearing three copies of chrXV. Trisomic chromosomes can form various synaptic configurations such as a fully synapsed pair plus a completely unsynapsed chromosome (“II+I”), synaptic partner switches, or triple synapsis along the entire chromosome length (Goldstein 1984; Loidl 1995; Robles et al. 2013). In “II+I” or synaptic partner switch configurations, at least one of the three homologs lacks a partner at every position along the chromosome length (**Fig. 1A**).

### More DSBs for longer times on homeologous and monosomic chromosomes

If homolog engagement functions chromosome-autonomously *in cis*, the homeologous and monosomic chromosomes should experience prolonged and more DSB formation compared to engagement-proficient chromosomes. To test this, we measured DSBs across meiotic time courses by separating high molecular weight DNA using pulsed-field gel electrophoresis (PFGE) followed by Southern blotting and indirect end-labeling (**Fig. 1B**). For each culture, we quantified DSBs on chrV (wild type, homeologous, or monosomic) and on chrIII as an internal control (**Fig. 1C**). To standardize quantitative comparisons despite different chrV copy numbers, we probed for a *natMX4* cassette integrated near the right end of one (or the only) *S. cerevisiae* copy of chrV. ChrIII was detected using a probe for *CHA1* on the left arm. DSB frequencies (broken DNA molecules as percent of total DNA) were corrected for multiple breaks on the same chromatid using a Poisson approximation (Murakami and Keeney 2014). In wild type, chrIII and chrV coincidentally formed DSBs with indistinguishable amounts and timing (**Fig. 1C**). By contrast, the homeologous and monosomic strains reproducibly displayed higher levels of broken chrV compared to chrIII. Similar increases were apparent (~1.2-fold for homeologous, ~1.7-fold for monosomic) whether considering peak values or time-averaged DSBs (area under the curve) (**Fig. 1C, D and Supplemental Fig. S2B**). The greater effect for monosomic chrV than for homeologous may reflect a difference in degrees of homolog engagement defect, because the homeologous chromosome pair still has some level of sequence similarity, whereas the monosomic chromosome completely lacks a pairing partner.

We also noted that the homeologous strain had similar DSB amounts on chrV and chrIII at the earliest time points (2 and 3 h), with chrV diverging from chrIII later (**Fig. 1C**). Moreover, we observed a reproducible delay of ~15 min on homeologous chrV compared to wild type when estimating DSB peak signal times by curve fitting (**Supplemental Fig. S2B,C**). (Absence of 3-h time points precluded this analysis for the monosomic cultures).

These results are consistent with the prediction that homeologous and monosomic chromosomes should accumulate more DSBs because of prolonged DSB formation. However, an alternative interpretation could be that DSB numbers are the same but DSB lifespan has been increased because of a repair delay caused by absence of a homolog as a repair partner (de Massy et al. 1994; Hochwagen et al. 2005). We did not favor this alternative as the sole explanation because repair using the sister chromatid can be rapid and efficient if a homolog is not available (Goldfarb and Lichten 2010). Nevertheless, we addressed this question more directly using an orthogonal approach to quantifying relative DSB formation: Spo11-oligonucleotide (oligo) sequencing.

### A homeologous chromosome pair incurs more DSBs

Spo11 cleaves DNA in a topoisomerase-like manner, creating a covalent bond at the 5′-strand termini of DSBs that is then clipped endonucleolytically to release Spo11 still attached to a short oligo (Neale et al. 2005) (**Supplemental Fig. S3A**). Spo11-oligo complexes are a quantitative byproduct of DSB formation and their lifespan is not tied to that of DSBs (Lange et al. 2011; Thacker et al. 2014). We immunoprecipitated Flag-tagged Spo11 from the homeologous strain harvested at 4 and 5 h in meiosis, then purified Spo11 oligos to prepare libraries for Illumina sequencing and compared with existing wild-type maps generated using a protein A-tagged version of Spo11 (Thacker et al. 2014; Mohibullah and Keeney 2017) (**Supplemental Fig. S3A**). Spo11-oligo maps from the wild-type and homeologous strains agreed well on chrIII (an internal control), exhibiting peaks (hotspots) with similar distributions at kilobase size scales (**Supplemental Fig. S3B**).

To compare per-chromosome distributions of Spo11 oligos, we first normalized the total number of sequence reads from each data set to one million (reads per million, RPM). If a specific chromosome (namely, chrV) generated more Spo11-oligo complexes, RPM on the other chromosomes would decrease even if the actual number of Spo11 oligos generated *in vivo* was unchanged. To account for this, we further scaled each data set to have an equal number of total RPM coming from the 15 chromosomes other than chrV. The scaled RPM was then summed for each chromosome. For the homeologous strain, we summed reads for both *S. pastorianus* and *S. cerevisiae* chrV.

As expected, the 15 chromosomes other than chrV aligned well with the diagonal when wild type and the homeologous strain were compared (gray points in **Fig. 2A**). This indicates that the relative number of Spo11-oligo reads between chromosomes is reproducible if they have a homologous partner. ChrV also fell on the diagonal at 4 h, but it deviated substantially at 5 h, with a 1.7-fold higher number of Spo11 oligos than expected from behavior of the homologous pair in wild type (orange points in **Fig. 2A, B**).

**Figure 2.**
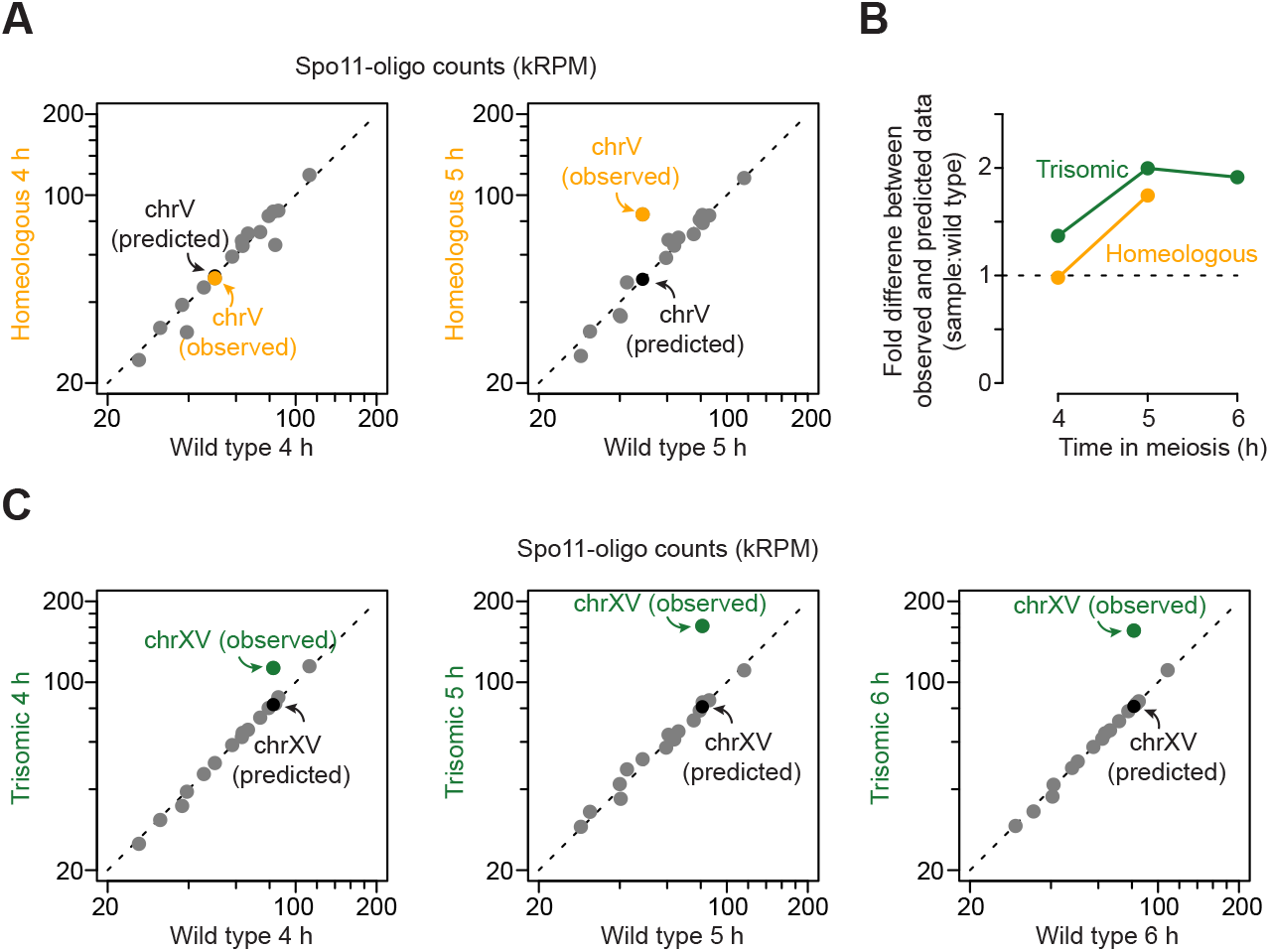
Increased DSBs are observed specifically on homeologous chrV and trisomic chrXV. (A, C) Comparison of per-chromosome Spo11-oligo totals (kRPM, thousands of reads per million reads mapped) between homeologous and wild type in A (left, 4 h; right, 5 h), and between trisomic and wild type in C (left, 4 h; middle, 5 h; right, 6 h). Each dot represents one chromosome. A predicted data point assuming that the homeologous chrV or trisomic chrXV has the same number of reads as the homologous chrV or chrXV is shown in each graph. (B) The fold difference between experimental and predicted values for the karyotypically abnormal strains (from A and C) at different meiotic times. Dashed line marks no change, meaning the experimental data matches prediction.

Because we are measuring normalized rather than absolute Spo11-oligo frequencies, we cannot exclude the possibility that the homeologous strain experiences changes in absolute DSB numbers on the other 15 chromosomes. However, such a change would have had to affect all of the chromosomes in close proportion to their DSB levels in wild type to maintain the good overall fit between the data sets (**Fig. 2A**). Therefore, these results strongly indicate that a homeologous chromosome pair selectively generates higher numbers of DSBs in a chromosome-autonomous fashion. The time dependence further supports the conclusion that homolog engagement defects allow DSBs to continue forming after they would normally have stopped.

### Trisomy also triggers elevated DSB formation

A further test of the homolog engagement model came from an accidentally generated aneuploid strain. In the course of other studies (Mohibullah and Keeney 2017), we observed that Spo11-oligo maps from a particular culture of a supposedly wild-type strain exhibited abnormally high read counts on chrXV. Other clones from the same stock behaved normally (Mohibullah and Keeney 2017), so we speculated that the exceptional culture might harbor a spontaneous aneuploidy for chrXV. Indeed, quantitative Southern blotting after PFGE revealed three copies of chrXV (**Supplemental Fig. S3C**). Attempts to obtain other aneuploid clones from the stock failed, so the trisomy-XV strain exists now only in its Spo11-oligo maps at 4, 5, and 6 h. These maps proved informative about the effects of aneuploidy on homolog engagement.

To compare per-chromosome distributions, we applied a two-step normalization similar to the one described above: RPM normalization followed by scaling to set the total number of reads from the 15 chromosomes other than chrXV equal between wild type and the trisomic strain. To correct for chromosome copy number, reads from trisomic chrXV were further scaled by a factor of 2/3. Even after correction, chrXV reads were overrepresented by two fold at 5 h (**Fig. 2C**). Again, overrepresentation was time dependent in that Spo11-oligo counts were only ~1.4-fold higher at 4 h (**Fig. 2B, C**). Counts remained ~two-fold elevated at 6 h; we interpret the lack of further increase from 5 h to 6 h as a consequence of Ndt80-driven exit from prophase I and concomitant downregulation of DSB formation (Allers and Lichten 2001; Gray et al. 2013; Thacker et al. 2014). In other words, prophase I exit ends the window of opportunity during which homolog engagement defects can allow DSBs to accumulate.

### Increased DSB formation is not due to appearance of new hotspots

The increase in Spo11 oligos on homeologous chrV or trisomic chrXV could reflect appearance of additional DSB hotspots or an increase of DSBs in the same hotspots. To differentiate between these possibilities, we called hotspots using an algorithm that identifies sites with Spo11-oligo counts above a threshold of 2.3 times the genome average (Pan et al. 2011). This identified 222 hotspots on *S. cerevisiae* chrV in the homeologous strain and 185 in wild type (**Fig. 3A**). However, the 37 hotspots called uniquely in the homeologous strain were weak ones that in wild type also had clusters of Spo11-oligo reads below the hotspot-calling threshold (**Fig. 3B, C**). Thus, the apparent increase in hotspot number on chrV when it is combined with a homeologous partner is a consequence of applying an arbitrary hotspot-calling threshold when there is a general increase in DSBs specifically on that chromosome. A similar conclusion was drawn for trisomic chrXV (**Fig. 3D–F**). We conclude that homolog engagement defects lead to increased DSB formation principally within existing hotspots rather than creating new sites of preferential Spo11 action.

**Figure 3.**
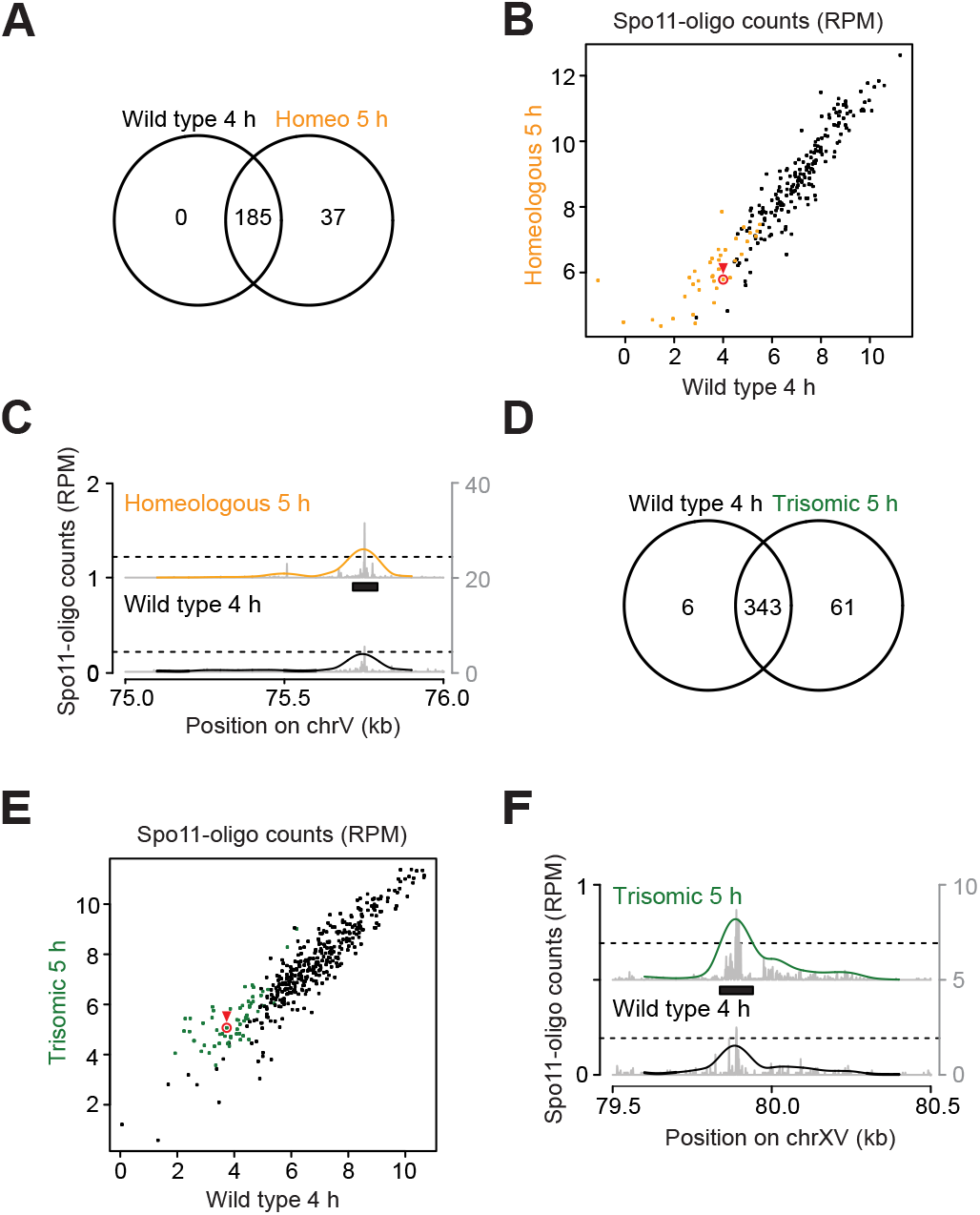
Conserved hotspots on homolog-engagement defective chromosomes. (A, D) Venn diagrams show the degree of overlap between hotspots called on chrV in wild type and the homeologous strain (A) or on chrXV in wild type and the trisomy-XV strain (D). (B, C, E, F) Comparison of hotspot strengths. Summed Spo11-oligo read counts are shown for all hotspots called on chrV (B) and chrXV (E). Orange dots are hotspots called only in the homeologous strain and green dots are those called only in the trisomic strain. Example hotspots are highlighted in red circle and arrowhead, and shown in (C, F). Black bars, boundaries of the called hotspots; line profiles, smoothed with a 201-bp Hann window; left Y-axis, RPM of smoothed profile; right Y-axis, RPM of raw data shown in gray.

### Homeology does not further increase DSB formation in a zip3 background

If the increased DSB formation on homeologous chromosomes reflects loss of the same feedback loop that is defective in ZMM mutants, as we hypothesized, then a ZMM mutation should be epistatic with homeology. That is, the global homolog engagement defect in a ZMM mutant would mean that a homeologous chromosome pair would not behave differently from homologous pairs.

To test this prediction, we generated Spo11-oligo maps from a *zip3Δ* strain carrying the homeologous chrV pair, which showed meiotic arrest similar to *zip3Δ* (**Supplemental Fig. S2A**). Homologous and homeologous chrV pairs exhibited similar numbers of Spo11-oligo reads in the *zip3Δ* background, with per-chromosome read counts from all 16 chromosomes aligning well on a diagonal (**Fig. 4A**). Because Spo11 oligos are normalized as RPM, this result shows that the per-chromosome DSB number as a fraction of total DSBs is the same in both strains for each chromosome pair. Since there is a cell-wide increase in DSBs in the *zip3Δ* mutant (Thacker et al. 2014), this further implies that the elimination of DSB overrepresentation on homeologous chrV was not due to lower break formation on homeologous chrV in *zip3Δ*, but instead to a genome-wide defect that caused all chromosomes to experience increased DSBs compared to *ZIP3*, irrespective of homology or homeology.

**Figure 4.**
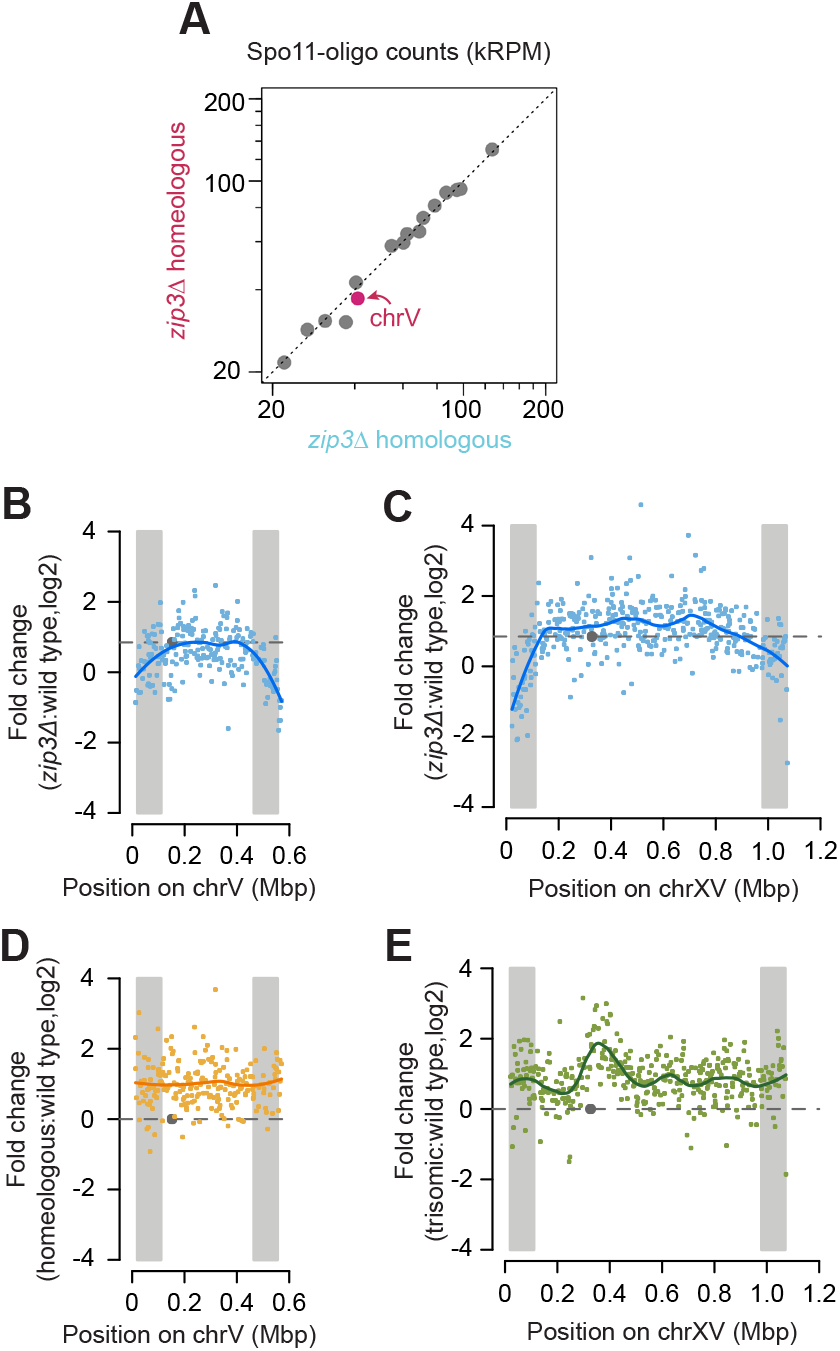
Loss of DSB overrepresentation on homeologous chromosomes in a zip3 mutant background. (A) Comparison of per-chromosome Spo11-oligo totals between *zip3Δ* homeologous and *zip3Δ* homologous maps. (B–E) Fold change of Spo11-oligo reads in hotspots along chrV (B, D) and chrXV (C, E) between different strains versus wild type as labeled. For the purpose of chromosome copy number correction, the reads on homeologous chrV were doubled and the reads on trisomic chrXV were multiplied by 2/3. For the *zip3Δ* map a scaling factor of 1.8-fold was applied to account for the global increase in the number of Spo11-oligo complexes in this strain (Thacker et al. 2014). Lines, local regression (loess); dashed horizontal lines, no change for (D, E) and genome average change for (B, C); gray circles, centromeres; gray shading, EARs (defined as the regions from 20 to 110 kb from telomeres).

### Effects of homeology and trisomy on within-chromosome DSB patterns

Homolog engagement shapes the DSB landscape because certain sub-chromosomal domains respond differently to the *zip3Δ* mutation: DSBs are increased less than the genome average in regions close to telomeres (within ~20 kb), around centromeres, and flanking the ribosomal DNA (rDNA) (Thacker et al. 2014). Moreover, chromosome end-adjacent regions (EARs, from ~20 to ~110 kb from telomeres) tend to be less sensitive to DSB suppression by homolog engagement, so on average they continue to experience DSB formation later into prophase I than is typically seen for interstitial regions (Subramanian et al. 2019).

To display regional responses to loss of feedback from homolog engagement, we plotted the ratio of Spo11-oligo counts in *zip3Δ* relative to wild type within each hotspot along the lengths of chrV and chrXV (**Fig. 4B,C**). As expected (Subramanian et al. 2019), the EARs (shaded in gray) showed lower ratios than did interstitial parts of these chromosomes (**Fig. 4B,C**). In other words, because EARs are less suppressed by homolog engagement in wild type, they display less of a DSB increase in *zip3Δ*. Interestingly, however, we did not observe this distinct behavior for EARs when plotting ratios of homeologous chrV (**Fig. 4D**) or trisomic chrXV (**Fig. 4E**) relative to wild type. In addition, trisomic chrXV showed a disproportionately large increase in Spo11 oligos emanating from an ~225 kb region near the centromere (**Fig. 4E**), not seen in *zip3Δ* (**Fig. 4C**). Mechanisms that may account for the different responses of these chromosomes to aneuploidy as opposed to absence of Zip3 are addressed below (Discussion). Regardless of the cause of the difference, however, these results show that the specific nature of a homolog engagement defect can shape how the DSB landscape changes, in turn emphasizing the importance of reactive feedback control mechanisms in molding DSB distributions within chromosomes.

### Homolog engagement displaces Rec114

To test the hypothesis that feedback from homolog engagement works through loss of DSB-promoting factors from chromosomes, we measured binding of myc-tagged Rec114 to homologous and homeologous chrV by chromatin immunoprecipitation followed by quantitative PCR (ChIP-qPCR). ChIP efficiencies (percent of input) were measured across meiotic time courses using five primer pairs targeting previously defined Rec114 ChIP peaks on *S. cerevisiae* chrV and on chrIII and chrVI as internal controls (**Fig. 5A, Supplemental Fig. S4A and Supplemental Table S2**) (Murakami and Keeney 2014).

**Figure 5.**
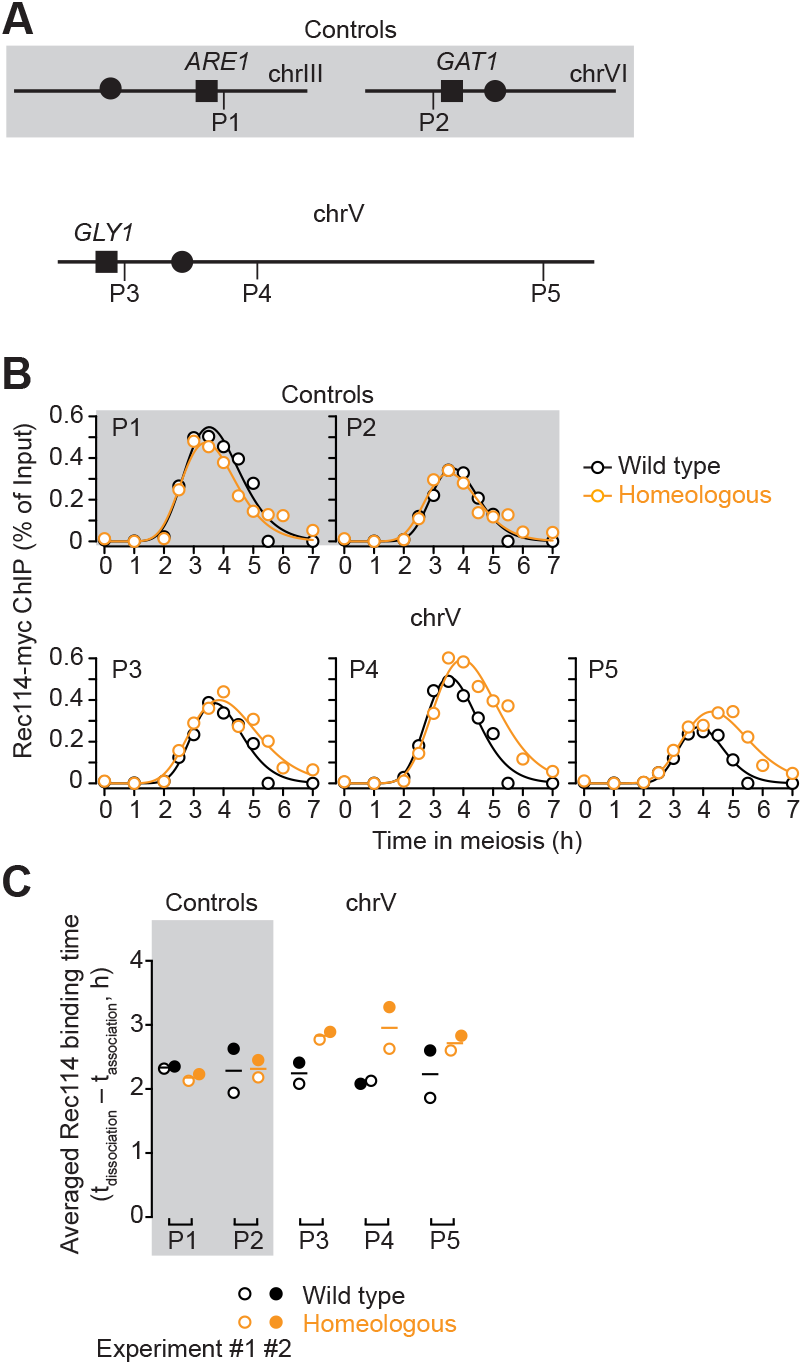
Persistent binding of Rec114 on homeologous chrV. (A) Primer pairs used for Rec114 ChIP-qPCR. Not to scale. P1 (on right arm of chrIII) and P2 (on left arm of chrVI) are internal controls (gray shading). P3 (left arm), P4 and P5 (right arm) are on chrV (no shading). (B) Rec114 binding kinetics for all five primer sets (data set #1). (C) Averaged Rec114 binding duration for all five primer sets from two data sets. The binding duration is defined as dissociation time minus association time. Means are indicated by horizontal lines. Data set #1 is shown as open circles and data set #2 is shown as filled circles.

In wild type, Rec114 ChIP showed similar kinetics at all five loci assayed, with interpolated peak times around 3.5 h (**Fig. 5B**). The homeologous strain also showed similar binding profiles for the two control loci, as expected, but the three loci on homeologous chrV continued to accumulate Rec114 beyond the time when levels started to decline in wild type, reaching higher peaks at later times (**Fig. 5B**). As a result, the duration of Rec114 binding was lengthened (**Fig. 5C and Supplemental Fig. S4B**). These results support the interpretation that defects in homolog engagement result in chromosomes spending more time in a DSB-competent state.

### SC formation is essential for DSB suppression by homolog engagement

Formation of both SC and crossovers is defective in ZMM mutants (Lynn et al. 2007; Pyatnitskaya et al. 2019), so these mutants are uninformative about which of these processes is the “homolog engagement” that establishes feedback control of DSB formation. To address this question, we turned to *gmc2Δ* and *ecm11Δ* mutations, which separate crossing over from SC formation (Humphryes et al. 2013; Voelkel-Meiman et al. 2016). Gmc2 and Ecm11 are components of the SC central element and function as a complex in facilitating the polymerization of the transverse filament protein Zip1 (Comino et al. 2001; Zavec et al. 2004; Humphryes et al. 2013). Either deletion leads to SC assembly defects, but meiotic divisions are completed efficiently with only modest delay (**Supplemental Fig. S5A**) and interhomolog crossovers form at elevated levels (1.1 to 2.8-fold higher than wild type, depending on the genetic interval assayed) (Voelkel-Meiman et al. 2016). We tested if these mutants exhibit signatures of homolog engagement defects.

First, we examined global DSB levels by quantifying Spo11-oligo complexes. Flag-tagged Spo11 was immunoprecipitated from meiotic cell extracts and the 3′ ends of Spo11 oligos were radiolabeled with terminal deoxynucleotidyl transferase and [α-^32P^]-dCTP before separation on SDSPAGE (**Fig. 6A, B and Supplemental Fig. S3A**). Both mutants generated substantially higher levels of Spo11-oligo complexes than wild-type cultures processed in parallel, with *gmc2Δ* reaching a peak level that was 1.8-fold higher than wild type and *ecm11Δ* reaching 1.7-fold higher (**Fig. 6A, B**). These increases are comparable to those for ZMM mutants *zip3Δ*, *zip1Δ*, and *msh5Δ* (Thacker et al. 2014). The increased numbers of DSBs in *ecm11Δ* and *gmc2Δ* presumably explain much (and possibly all) of the elevated crossing over (Voelkel-Meiman et al. 2016).

**Figure 6.**
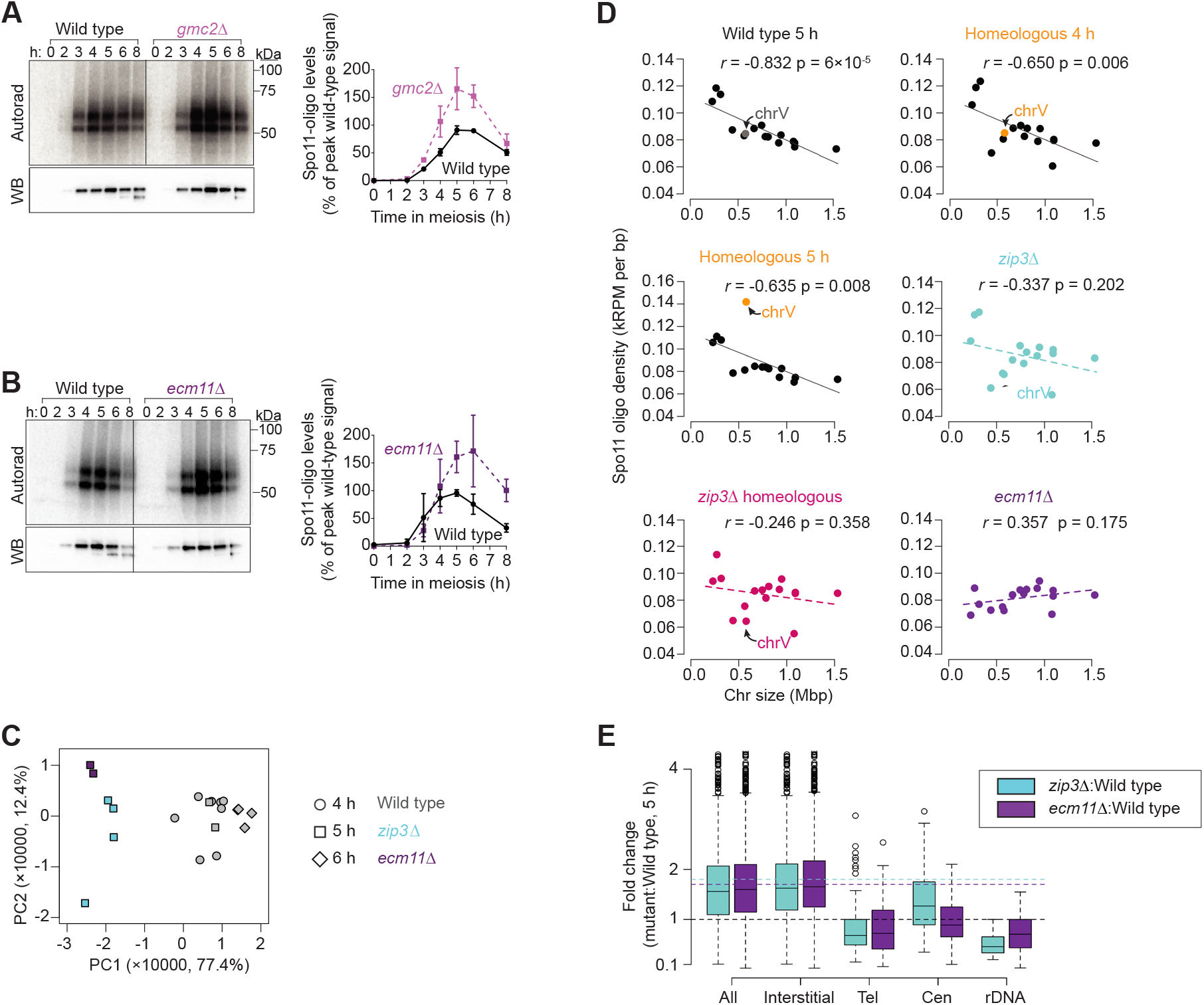
*gmc2Δ* and *ecm11Δ* mutants share similar homolog-engagement signatures as zip3Δ. (A, B) Representative labeling of Spo11-oligo complexes in *gmc2Δ* (A) and *ecm11Δ* (B) mutants and quantification relative to wild-type cultures processed in parallel. Error bars indicate mean ± SD for 3 cultures. Radiolabeled Spo11-oligo complexes are detected by autoradiography (left panel, top) and total Spo11 is detected by anti-Flag western blot (WB; left panel, bottom). The two main labeled species differ in the sizes of oligos (Neale et al. 2005). Most Spo11 protein does not end up making DSBs, so nearly all of the visible western blot signal is from free Spo11 that does not have an oligo attached (Neale et al. 2005). (C) Principal component analysis of 21 wild-type and mutant Spo11-oligo maps. (D) Loss of anticorrelation between chromosome length and DSB density in homolog-engagement-defective mutants. Each point is one chromosome. Correlation coefficients (Pearson’s r) are shown. (E) Fold change of Spo11-oligo counts in different chromosomal domains. Tel, within 20 kb of telomeres; Cen, within 10 kb of centromeres; rDNA, from 60 kb leftward to 30 kb rightward of rDNA; Interstitial, all others. Horizontal dashed lines mark values assumed as no change (black) and average change (1.8-fold for *zip3Δ*, cyan; 1.7-fold for *ecm11Δ*, purple). Boxes indicate median and interquartile range; whiskers indicate the most extreme data points that are ≤ 1.5 times the interquartile range from the box; individual points are outliers

Next, we compared *ecm11Δ* and *zip3Δ* Spo11-oligo maps. To evaluate similarities and differences systematically, we applied principal component analysis to multiple wild-type and mutant Spo11-oligo maps generated in this study and previous ones (Thacker et al. 2014; Zhu and Keeney 2015; Mohibullah and Keeney 2017; Murakami et al. 2020) (**Fig. 6C**). The first and second principal components (PC1 and PC2) together accounted for 89.8% of the variance among the data sets. PC1 separated both *ecm11Δ* and *zip3Δ* mutants from wild type, indicating that these mutants share DSB landscape features in common with one another. This conclusion was reinforced by hierarchical clustering, which grouped *ecm11Δ* and *zip3Δ* datasets with one another separately from wild type (**Supplemental Fig. S5B**).

Per-chromosome Spo11-oligo densities are negatively correlated with chromosome length in wild type, and this correlation collapses in a *zip3Δ* mutant (**Fig. 6D**) (Thacker et al. 2014). Loss of chromosome-size-dependent regulation of DSB numbers is thus a hallmark of defects in homolog engagement (Keeney et al. 2014; Thacker et al. 2014; Subramanian et al. 2019; Murakami et al. 2020). The *ecm11Δ* mutant showed this same hallmark (**Fig. 6D**). In contrast, presence of one homeologous pair should not eliminate the negative correlation of DSB density with chromosome size for the rest of the genome, because homolog engagement functions chromosome autonomously. Indeed, the homeologous strain retained the overall negative correlation except for chrV (**Fig. 6D**).

Spo11-oligo maps in *ecm11Δ* also displayed patterns similar to *zip3Δ* within specific sub-chromosomal domains: Spo11-oligo levels were increased less than genome average in regions near telomeres, centromeres, and the rDNA array on chrXII (**Fig. 6E and Supplemental Fig. S5C–E**).

We conclude that *ecm11Δ* (and by extension, *gmc2Δ*) phenocopies multiple signatures of the DSB dysregulation previously documented in *zip3Δ.* Because *ecm11Δ* and *gmc2Δ* mutants are SC-deficient but crossover-proficient, these results strongly indicate that SC formation is essential for DSB regulation by homolog engagement and that crossover formation in the absence of SC is not sufficient to down-regulate DSB formation.

## Discussion

We have shown that individual yeast chromosomes unable to engage with a homologous partner continue to accumulate DSBs past the time when other (homolog-engagement-proficient) chromosomes in the same cell have largely stopped breaking. By examining crossover-proficient mutants that are unable to make mature SC because they lack key components of the central element, we also provide evidence that synapsis per se is required for homolog engagement-mediated DSB suppression. Kim, Shinohara and colleagues have provided independent and distinct evidence for increased DSB formation in *gmc2* and *ecm11* mutants (Lee et al. 2020); our results agree well with theirs. These findings support the conclusion that feedback control of DSB formation in yeast works in a chromosome-autonomous fashion as a response to SC formation, consistent with cytological evidence in mouse (Wojtasz et al. 2009; Kauppi et al. 2013) but apparently distinct from nucleus-wide responses to crossover defects in *C. elegans* and possibly *D. melanogaster* (Bhagat et al. 2004; Carlton et al. 2006; Joyce and McKim 2010; Joyce and McKim 2011; Rosu et al. 2013; Stamper et al. 2013; Keeney et al. 2014; Crown et al. 2018).

One key difference between these species is that DSB formation precedes and is required for homologous synapsis in yeast and mouse (Alani et al. 1990; Padmore et al. 1991; Baudat et al. 2000; Romanienko and Camerini-Otero 2000; Mahadevaiah et al. 2001), whereas DSB formation usually occurs after SC formation and is dispensable for homologous synapsis in *C. elegans* and *D. melanogaster* (Dernburg et al. 1998; McKim et al. 1998; Alpi et al. 2003; Colaiacovo et al. 2003). Thus, the presence of SC can serve for cells to sense whether DSB formation has served its purpose in yeast and mouse, but would not be informative for this in *C. elegans* and *D. melanogaster*. These findings illustrate how evolutionarily distinct strategies for completing the meiotic program can place different constraints on the mechanisms available for cells to ensure that DSBs form where they are needed but stop being formed where they have already done their job.

Our findings further support the interpretation that homolog engagement works by promoting the dissociation of DSB-promoting factors from chromosomes after SC formation (Wojtasz et al. 2009; Carballo et al. 2013; Keeney et al. 2014; Subramanian et al. 2019; Murakami et al. 2020). In mouse, axis proteins HORMAD1 and HORMAD2 are displaced from all synapsed axes, dependent on the AAA+ ATPase TRIP13 (Wojtasz et al. 2009; Roig et al. 2010). Synapsis also displaces specialized assemblies of SPO11-accessory proteins REC114 and MEI4 that form on the pseudoautosomal region, the only part of the X and Y chromosomes that recombines in male meiosis (Acquaviva et al. 2020). In yeast, the TRIP13 ortholog Pch2 also directs relocalization of Hop1 (ortholog of HORMAD1) after SC formation (Borner et al. 2008; Joshi et al. 2009; Subramanian et al. 2016; Raina and Vader 2020). Moreover, many proteins that promote Spo11 activity (Mei4, Rec102, Rec104, Rec114, and Red1) all dissociate from synapsed chromosomes (Smith and Roeder 1997; Kee et al. 2004; Li et al. 2006; Maleki et al. 2007; Panizza et al. 2011; Carballo et al. 2013). A plausible scenario is that Pch2 is recruited or activated in the context of SC and disrupts the interaction between the HORMA domain in Hop1 and its direct binding partner, the “closure motif” in Red1 (Chen et al. 2014; Kim et al. 2014; West et al. 2018). This locally terminates the DSB competent state by dissociating or remodeling Hop1 concomitant with dissociation of DSB-promoting proteins.

While our study argues that crossing over in the absence of SC is not sufficient in yeast to provoke DSB suppression via homolog engagement, we cannot rule out the possibility that crossover formation might cooperate with SC and play some role in regulating DSB formation. Although this question has not been directly addressed in mouse, either, it is noteworthy that the SC that forms between nonhomologous chromosomes in the absence of recombination in a *Spo11^−/−^* mutant is sufficient to provoke displacement of HORMAD1 and HORMAD2 (Wojtasz et al. 2009) and chromosome axis remodeling plus REC114 displacement from the pseudoautosomal region (Acquaviva et al. 2020). This is consistent with the possibility that synapsis is both necessary and sufficient to downregulate DSB formation.

TRIP13-dependent displacement of HORMAD1 and HORMAD2 in mouse and Rec102 and Rec104 displacement in yeast occur with some delay after SC formation (Kee et al. 2004; Wojtasz et al. 2009), implying that downregulation of DSB formation is also not instantaneous upon completion of synapsis. This delay might simply reflect the time necessary for TRIP13/Pch2 to be recruited and to work, or it might indicate that some time-dependent change in the SC is needed as the signal for TRIP13/ Pch2 to act. In this context, we note that progression of recombination toward crossover formation is associated with pronounced changes in SC structure in *C. elegans* (Libuda et al. 2013; Pattabiraman et al. 2017; Rog et al. 2017; Woglar and Villeneuve 2018; Zhang et al. 2018; Cahoon et al. 2019). A view that might unify DSB control systems in *C. elegans*, yeast, and mouse might be that “maturation” of SC structure (tied to crossing over in *C. elegans* but perhaps tied only to the act of synapsis itself in yeast and mouse) is a conserved signal for local downregulation of DSB formation.

Different chromosomal subdomains respond differently to the defect in DSB regulation in ZMM mutants (Thacker et al. 2014). To our surprise, even though *zip3* mutation was epistatic with homeology, we found that the karyotypic abnormalities we examined did not cause the same subchromosomal changes as a *zip3* mutation. We envision two nonexclusive possibilities to account for the different responses of these chromosomes to aneuploidy as opposed to absence of Zip3. First, these aneuploidies may cause less severe homolog engagement defects along the entire chromosome lengths than *zip3* mutation does. In this scenario, the aneuploid chromosomes retain some residual DSB suppression, so interstitial regions experience less of an increase in DSB formation compared to *zip3Δ*. Second, interstitial regions on the aneuploid chromosomes may retain more residual homolog engagement compared to EARs, for example if there is still some degree of interstitial SC formation in a fraction of cells in the population. Zip1 mediates homology-independent coupling of centromeres during prophase I (Kemp et al. 2004; Tsubouchi and Roeder 2005; Newnham et al. 2010), so it is possible that the homeologous centromeres might pair and sometimes nucleate SC. Moreover, although SC is restricted to homologous pairs in normal diploids, other synaptic configurations can occur, for example between homologous or nonhomologous chromosome segments in haploid, triploid, tetraploid, and hybrid yeast strains (Gillies et al. 1974; Moens and Ashton 1985; Loidl et al. 1991; Loidl 1995; Lorenz et al. 2002). In the majority of these cases, partial SC is produced along with unsynapsed axes and partner switches. Therefore, partial SC formed between the homeologous pair could possibly activate homolog engagement feedback to some extent.

A striking finding was the high degree of DSB overrepresentation around the centromere on trisomic chrXV. In fully triploid strains, triple-synapsed trivalents, “II+I” synapsis, and trivalents with partner switches have been observed (Loidl 1995), but the synaptic configuration of a single trisomic chromosome set is not known. We speculate that the pericentromeric region might be frequently left unsynapsed in trivalent configurations. Regardless of the cause, however, our findings suggest that trisomy antagonizes operation of the pathways that normally function to suppress DSB formation and crossing over near centromeres (Rockmill et al. 2006; Vincenten et al. 2015).

Although we have focused on the increase in DSB formation that accompanies defects in homolog engagement, it is important to emphasize that engagement-defective chromosomes do not simply continue to make more and more DSBs indefinitely. Other systems also come into play to restrict the amount of DSB formation that can occur, including feedback dependent on activation of the DSB-responsive kinase Tel1 (ATM in mammals) (Joyce et al. 2011; Lange et al. 2011; Zhang et al. 2011; Carballo et al. 2013; Anderson et al. 2015; Garcia et al. 2015) and a global shutdown in DSB formation tied to exit from the pachytene stage driven by the Ndt80 transcription factor (Allers and Lichten 2001; Gray et al. 2013; Thacker et al. 2014). The complex interplay between multiple DSB regulatory pathways helps to explain how the meiotic program can be robust in the face of whole chromosome aneuploidy as examined here, and possibly in similar situations in humans such as Down syndrome (trisomy 21), Turner syndrome (45, XO) and Klinefelter syndrome (47, XXY). These considerations likely also apply to challenges posed by heterozygosity for translocations or large chromosomal deletions, insertions, duplications, or inversions and by decreased homology between chromosomes in outcrosses. Our findings thus illustrate basic principles that contribute to the fidelity of the meiotic program.

## Materials and Methods

### Yeast strains

Yeast strains used in this study are of the SK1 background (**Supplemental Table S3**). *SPO11* was C-terminally tagged with three Flag epitope repeats by targeted integration of a *6His-3FLAG-loxP-kanMX-loxP* construct amplified from an *S. cerevisiae* SK1 *SPO11-FLAG* strain provided by Kunihiro Ohta, Univ. Tokyo (Kugou and Ohta 2009) in the following strains: homeologous strain, monosomic strain, *zip3Δ* mutant*, zip3Δ* homeologous mutant, *gmc2Δ* and *ecm11Δ* mutant. In the trisomic strain, Spo11 was protein-A (PrA) tagged (Thacker et al. 2014). We also used published Spo11 oligo maps generated using Flag-tagged Spo11 in wild type (Zhu and Keeney 2015; Murakami et al. 2020) and PrA-tagged Spo11 in wild type and *zip3Δ* strains (Thacker et al. 2014; Mohibullah and Keeney 2017). The nourseothricin drug resistant marker, *natMX4*, was inserted near the end of the right arm of chrV (the convergent intergenic region between *PUG1* and *YER186c*: corresponding to coordinates 560580-1 of S288C genome assembly from SGD (*Saccharomyces* Genome Database)) in a wild-type SK1 haploid strain. Yeasts were transformed using standard lithium acetate methods. Correct tagging was verified by PCR and Southern blot. The *zip3*, *gmc2* and *ecm11* deletion mutants were generated by replacing the coding sequences with the hygromycin B drug resistance cassette (*hphMX4*) individually through yeast transformation. Gene disruption was verified by Southern blot.

The homeologous strain contains one copy of *S. pastorianus* chrV in an otherwise SK1 background. *S. pastorianus* (lager-brewing yeast) is a hybrid of *S. cerevisiae* and *S. eubayanus* (Gallone et al. 2018; Monerawela and Bond 2018). The copy of chrV introgressed from *S. pastorianus* into *S. cerevisiae* is originally derived from *S. eubayanus*. An *S. cerevisiae* SK1 haploid strain with chrV replaced by *S. pastorianus* chrV (marked with *ilv1*) was a gift from M. Lichten (Goldman and Lichten 2000). This strain was crossed with an SK1 haploid strain (chrV marked with *natMX4*) to create the homeologous diploid strain immediately before preparing premeiotic and meiotic cultures.

The trisomic chrXV strain arose spontaneously. A single clone obtained from the frozen stock of a wild-type strain was expanded and cultured in pre-sporulation and sporulation media, and samples were collected for Spo11-oligo mapping and PFGE as described previously (Mohibullah and Keeney 2017) and below. The karyotype was evaluated by quantitative Southern blotting of the PFGE samples using *GIT1* (chrIII) and *ARG1* (chrXV) open reading frames as probes. Other independent clones obtained contemporaneously from the same frozen stock were euploid, and we later attempted unsuccessfully to isolate additional trisomic clones from this stock.

The monosomic chrV strain was generated as follows (**Supplemental Fig. S1**). A strong *GAL1* promoter, marked with the *Kluyveromyces lactis URA3* gene, was inserted adjacent to centromere DNA to create a conditional centromere (Hill and Bloom 1987). Plasmid pCEN05-UG containing a chrV centromere-destabilizing cassette was provided by R. Rothstein (Reid et al. 2008). The *CEN5* targeting fragment was liberated by NotI digestion and transformed into an SK1 haploid to replace the native *CEN5*, confirmed by Southern blot. A previous study suggested that chromosome loss could be followed by endo-reduplication of the remaining chromosome, especially for small chromosomes, but chrV was not directly analyzed (Reid et al. 2008). A presporulation procedure was specifically developed to allow high frequency of chromosome loss (see Culture Methods), with loss of chrV and endo-reduplication events closely monitored by tetrad dissection and spore clone genotyping (**Supplemental Fig. S1 and Supplemental Table S1**).

### Culture methods

Synchronous meiotic cultures were prepared using the SPS pre-growth methods as described (Murakami et al. 2009) for all strains in various culture volumes required for different experimental purposes, except for the trisomic strain using the YPA pre-growth (Alani et al. 1990; Padmore et al. 1991) and for the monosomic chrV strain which is slightly modified and addressed in next paragraph. Two stages of SPS pre-growth culturing were performed to reproducibly obtain appropriate cell density before transferring into sporulation medium (SPM) for better synchrony. In brief, cells from a 4 ml saturated overnight YPD (1% yeast extract, 2% peptone, 2% glucose) were used to inoculate into 25 ml of SPS (0.5% yeast extract, 1% peptone, 0.67% yeast nitrogen base without amino acids, 1% potassium acetate, 0.05 M potassium biphtalate (pH 5.5), 0.002% antifoam 204 (Sigma)) to a density of 5 × 10^6^ cells/ml and cultured at 30°C at 250 rpm for 7 h. Cells were then inoculated into an appropriate larger volume of fresh SPS (200 ml for PFGE and labeling of Spo11-oligo complexes; 2 × 800 ml for Spo11-oligo mapping) at a density of 3 × 10^5^ cells/ml and cultured at 30°C at 250 rpm for 12–16 h until the density reached 3–4 × 10^7^ cells/ml. Cells were collected by centrifugation or filtration, washed with 2% potassium acetate, then resuspended at 4 × 10^7^ cells/ml in an appropriate volume of SPM (2% potassium acetate, 0.001% polypropylene glycol) (100 ml for PFGE and Spo11-oligo complexes labeling; 1 l for Spo11-oligo mapping) supplemented with 0.32% amino acids complementation medium (1.5% lysine, 2% histidine, 2% arginine, 1% leucine, 0.2% uracil, 1% tryptophan). Cultures were incubated in a 30°C shaker at 250 rpm to induce sporulation. Samples were collected at desired times after transferring into SPM.

One round of 13.5 h culture in YPA (1% yeast extract, 2% peptone, 1% potassium acetate) was used in the place of the SPS pre-growth for sporulating the trisomic strain.

The modifications for the monosomic strain were as follows. Single colonies were patched on YP-galactose plates for 24 h at 30°C, then to freshly made 5-FOA plates (containing 2 g/l 5-fluoroorotic acid) for 24 h at 30°C. Standard synchronous meiotic cultures were then prepared as described above. Samples were collected at 24 h for tetrad dissection. Complete loss of one copy of chrV should yield tetrads with 2:2 (live:dead) segregation, with all viable spores lacking the *Kl-URA3* marker. Endoreduplication of chrV would give four viable *ura3* spores. Failure to induce centromere loss with yield four viable spores with 2:2 segregation patterns for the *Kl-URA3* marker (**Supplemental Fig. S1**). Only cultures for which all tested tetrads had the pattern expected for complete loss and lack of endoreduplication were processed for downstream applications (**Supplemental Table S1**).

To check meiotic divisions, aliquots were collected at various times from synchronous meiotic culture, fixed in 50%(v/v) ethanol and stained with 0.05 μg/ml 4’, 6-diamidino-2-phenylindole (DAPI). Mono-, bi-, and tetranucleate cells were scored by fluorescence microscopy.

### Detection of meiotic DSBs by Southern blot

Genomic DNA was prepared in plugs of low-melting point agarose as described (Borde et al. 2000; Murakami et al. 2009) to avoid random shearing. The high-molecular-weight DNA was separated by PFGE as described (Borde et al. 2000; Murakami et al. 2009) and then probed by Southern blot using a radiolabeled DNA fragment within the *CHA1* coding sequence located near the left arm end of chrIII (internally controlled chromosome for PFGE) or the *natMX4* cassette on *S. cerevisiae* chrV. Signals were detected by phosphorimager, and quantified with ImageGauge software (Fujifilm). DSB frequencies at different time points were calculated as percentages of broken molecule signals divided by paternal signals plus broken molecule signals in each lane. Observed DSB frequencies were Poisson corrected as described before (Murakami and Keeney 2014) for correction of the situation that multiple DSBs happened on each chromosome.

### End-labeling Spo11-oligo complexes and Spo11-oligo mapping

Spo11-oligo complexes were extracted and detected as previously described (**Supplemental Fig. S3A**) (Neale and Keeney 2009). Briefly, Spo11-oligo complexes were immunoprecipitated from whole-cell extracts by using mouse monoclonal anti-Flag M2 antibody (Sigma). Precipitated Spo11-oligo complexes were end-labeled with [α-^32P^] dCTP in a terminal deoxynucleotidyl transferase reaction, resolved by SDS-PAGE, then transferred onto PVDF membrane and visualized by phosphorimager. Blots were probed with mouse monoclonal anti-Flag M2 conjugated to horseradish peroxidase (Sigma) and detected by chemiluminescent (ECL+ or ECL Prime, Amersham).

For Spo11-oligo mapping, sporulation cultures of different volumes (450 ml for homeologous strain; 300 ml for trisomic strain; 600 ml for all other strains) were harvested at desired time points after transferring to sporulation media. Maps were generated for this study in strains carrying *SPO11-Flag* as described (Lam et al. 2017) except for the trisomic strain carrying *SPO11-PrA*. Previously published wild-type maps (Thacker et al. 2014; Mohibullah and Keeney 2017) were used as controls in this study. For the PCA and clustering analyses, we also included other wild-type and *zip3Δ* maps (Zhu and Keeney 2015; Mohibullah and Keeney 2017; Murakami et al. 2020).

### ChIP for Rec114-Myc

Strains expressing Rec114 with C-terminal tag of 8 copies of the Myc epitope (*REC114-Myc*) were generated as described (Murakami and Keeney 2014). Tagged Rec114 was functional, as *REC114-Myc* and diploids showed normal spore viability.

Samples of 50 ml (2 × 109 cells) were collected at desired times after transferring to SPM (0, 1, 2, 2.5, 3, 3.5, 4, 4.5, 5, 5.5, 6, 7 h) and crosslinked with 1% formaldehyde for 30 min at room temperature. Crosslinking was terminated by incubation with 131 mM glycine for 5 min. Cells were washed twice with ice-cold TBS, frozen with liquid nitrogen and stored at −80°C. After re-suspending frozen cells using 1 ml of Lysis buffer (50 mM HEPES-KOH pH 7.5, 140 mM NaCl, 1 mM EDTA, 1% Triton X-100, 0.1% Na-deoxycholate, 1 mM PMSF, 7 μg/ml aprotinin, 1% protease inhibitor cocktail (Sigma), 1x Complete Protease Inhibitor Cocktail (Roche)) with ca. 900 μl of 0.5 mm zirconia/silica beads (BioSpec Products) in 2 ml screw-cap Eppendorf tubes, cells were then disrupted by vigorous shaking 1 min × 12 times with an intensity of 6.5 M/s in a FastPrep24 (MP Biomedicals) to reach a cell breakage efficiency over 99% (99.3% in our experiment). Another 1 ml of Lysis buffer was added after cell disruption. Chromatin DNA in the whole cell extracts (WCE) was sheared by sonication with “M” intensity, 30 s ON/30 s OFF for 15 min × 3 times in Bioruptor Sonication System UCD200 (Diagenode) in 15 ml polystyrene conical tubes. Insoluble cell debris was removed by centrifugation at 21,130 *g*, 5 min, 4°C. WCE was further sonicated with the same condition one more time to yield average DNA size around 350 bp (range of 100 bp-500 bp). ChIP was performed as described (Murakami and Keeney 2014).

Locations of five primer pairs for quantitative PCR (qPCR) analysis are as followed: one locus on right arm of chrIII (P1), one locus on left arm of chrVI (P2, near strong DSB hotspot *GAT1*), and three loci on chrV (P3 to P5, P3 was near strong DSB hotspot *GLY1*) (Murakami and Keeney 2014) (primer sequences listed in **Supplemental Table S2**). qPCR was performed using the LightCycler^®^ 480 SYBR Green I Master (Roche) according to manufacturer recommendations. All measurements of ChIP and mock samples were expressed relative to the standard (dilution series of corresponding input samples).

### Bioinformatics analysis

#### Curve Fitting for DSB kinetic profile

To estimate the time of DSB formation, a log-normal curve defined below was fitted to the DSB frequency (% of lane) plotted as a function of time (h): y = a+b*exp(-(log(x+1)-c)^2/d^2 where x is time in hour, y is DSB frequency, a is the background, b is the peak height, c is the peak position, and d is the equivalent of standard deviation. We set the background parameter (a) to the DSB frequency at 0 h, then fitted the equation to the data points by least-squares to estimate the other parameters (b, c and d) using the “nls” function in R.

#### Spo11-oligo Mapping Analysis

Sequencing (Illumina HiSeq 2500, 2 x 50 bp paired-end reads) was performed by the MSKCC Integrated Genomics Operation. In silico clipping of library adapters and mapping to the genome was performed by the Bioinformatics Core Facility at MSKCC using a custom pipeline as described (Pan et al. 2011; Thacker et al. 2014) with modifications. A full copy of the source code is available online at http://cbio.mskcc.org/public/Thacker_ZMM_feedback. For strains that are pure *S. cerevisiae* SK1 background, Spo11-oligo reads were mapped to the sacCer2 genome assembly of type strain S288C from SGD (*Saccharomyces* Genome Database). For the homeologous chrV strain, the sequence of *S. eubayanus* chrV (GenBank accession number JMCK01000005.1) (Baker et al. 2015) was added as an extra chromosome to the customized pipeline. We used only the uniquely mapping reads. Analyses were performed using R version 3.4.0 or GraphPad Prism 7.0a.

Raw and processed sequence reads for new maps generated in this study (**Supplemental Table S4**) are deposited in the Gene Expression Omnibus (GEO) database (https://www.ncbi.nlm.nih.gov/geo/) (accession number GSE152957). This accession also contains the curated maps (unique mapping reads only, normalized to reads per million mapped) in wiggle format to allow direct visualization in appropriate genome browsers, e.g., the UCSC browser (https://genome.ucsc.edu/s) using genome version sacCer2. Previously published maps analyzed in this study are from GEO accession numbers GSE48299, GSE67910, GSE84696, GSE119689 (**Supplemental Table S5**).

Each map was normalized to the total number of reads that mapped uniquely to a chromosome (RPM; excluding reads mapping to rDNA, mitochondrial DNA, or the 2 micron plasmid). The chromosome copy number was corrected for the homeologous chrV pair by summing up the reads from of *S. cerevisiae* chrV copy and *S. pastorianus* chrV or by multiplying the *S. cerevisiae* chrV read count by a factor of two, as appropriate for the analysis. We corrected for chromosome copy number for trisomic chrXV by multiplying by 2/3. The purpose of this was to evaluate how much of the chrXV-specific increase in Spo11 oligos was in excess of the amount that might have been expected simply from the increase in chromosome copy number.

Since these copy-number corrections affect the total normalized read number, we made the following further adjustments for analytical purposes. For the homeologous strain, we calculated the total reads without chrV for both homeologous and wild type maps and averaged them to get a standard number. Then we adjusted the read counts for each chromosome including chrV using the standard number as total reads. We did the same separately for the trisomic strain.

For the analysis of DSB distributions within chrV in the homeologous strain (shown in **Fig. 4**), we could not sum the maps for the *S. cerevisiae* and *S. pastorianus* copies of chrV, so we instead doubled the reads from *S. cerevisiae* chrV. For *zip3Δ* and *ecm11Δ* maps, a scaling factor based on quantification of Spo11-oligo complexes was applied, as described in the appropriate figure legends.

DSB hotspots were defined as clusters of Spo11 oligos meeting cutoffs for cluster size and Spo11-oligo density as previously described (Pan et al. 2011). Briefly, candidate hotspots were first identified as chromosome segments where the Spo11-oligo map smoothed with a 201-bp Hann window was >0.193 RPM per bp, which is 2.3-fold over the genome average Spo11-oligo density. Adjacent hotspots separated by ≤ 200 bp were merged, then candidate hotspots were filtered to remove calls that were less than 25 bp wide and/or contained fewer than 10 RPM total.

### Principal Component Analysis and Hierarchical Cluster Analysis

In total, 20 maps were included in this study: 14 wild-type maps [two 4 h with Spo11-PrA (Thacker et al. 2014), two time course collected at 4, 5, 6 h with Spo11-PrA (Mohibullah and Keeney 2017), two 4 h with Spo11-flag (Zhu and Keeney 2015), two time course collected at 4 and 6 h with Spo11-flag (Murakami et al. 2020)], 4 *zip3Δ* maps [two 5 h with Spo11-flag in this study and two 5 h with Spo11-PrA (Thacker et al. 2014)] and 2 *ecm11Δ* maps with Spo11-flag (this study). The total number of Spo11-oligo read counts (normalized to RPM) were calculated on each. Principal component analysis on the per-chromosome total reads from 20 datasets was performed using the ‘princomp’ function in R. The first three principal components accounted for 77.4%, 12.4% and 2.5% of the variance across these data sets, respectively. Hierarchical clustering was performed with the ‘hclust’ function in R using Ward’s D2 method.

### Curve Fitting for Rec114 ChIP

The curve fitting method to define association and dissociation times was as described previously (Murakami and Keeney 2014). Briefly, a modified Gaussian curve was fitted to all qPCR data points to define the Rec114 ChIP signal peak position for each primer pair. Next, this peak was used to fit a saturating exponential growth (logistic) curve to just the upward slope of the ChIP profile. We defined the tassociation as the time point where the logistic curve reached 50% of the maximum. We also estimated the dissociation time of DSB protein by fitting a logistic curve to the down¬ward slope of the ChIP profile as tdissociation when the logistic curve reached 50% of the maximum

## Acknowledgments

We are grateful to Michael Lichten and Rodney Rothstein for providing strains and plasmids; Miki Shinohara and Keun Kim for discussions and sharing unpublished data; Dean Dawson and Amy MacQueen for sharing unpublished data; Agnes Viale (Memorial Sloan Kettering Cancer Center (MSKCC) Integrated Genomics Core Laboratory) for sequencing; Nicholas Socci (MSKCC Bioinformatics Core Facility) for mapping sequence reads; Stewart Shuman for gifts of T4 RNA ligase; and members of the S.K. laboratory, especially Shintaro Yamada and Devanshi Jain, for discussion and comments on the manuscript. MSKCC core facilities are supported by Cancer Center Support Grant P30 CA008748. This work was supported by NIGMS grant R35 GM118092 to S.K.

## Author contributions

X.M. performed experiments. X.M. analyzed Spo11-oligo map data with contributions from H.M. N.M. generated the Spo11-oligo map of the trisomy-XV strain. X.M., H.M., and S.K. designed the study and wrote the paper. H.M. and S.K. supervised the research. S.K. secured funding.

---

**Supplemental Figure S1.**
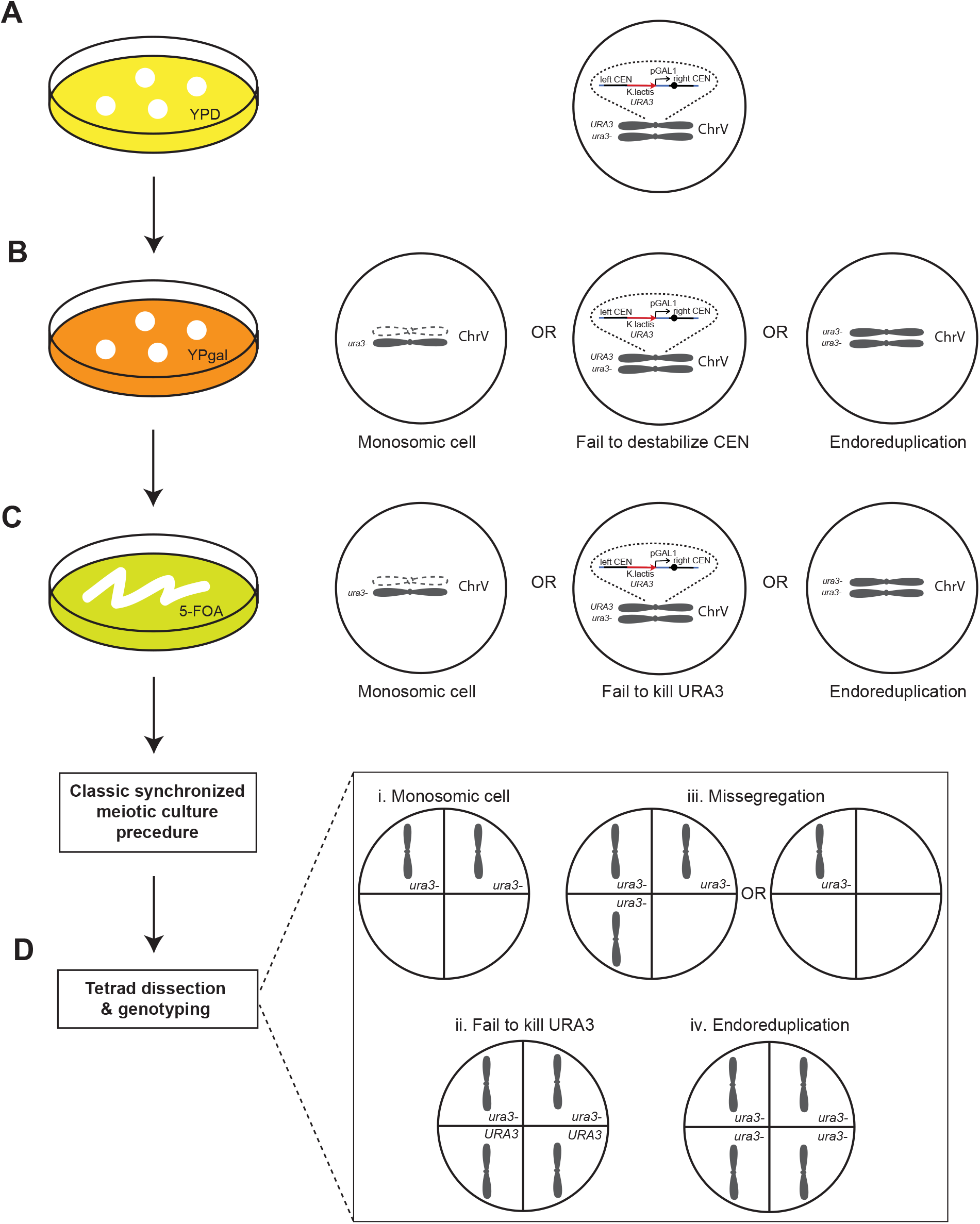
Strategy for constructing the monosomic strain. (A) In the diploid parental strain, one copy of chrV contains a conditional centromere generated by insertion of the GAL1 promoter (marked with URA3) adjacent to the centromere. Cells are cultivated on a YPD plate overnight before transferring to a YP galactose plate. (B) After 24 h galactose induction, most cells lose one copy of chrV due to the destabilized centromere. However, galactose induction may not be 100% efficient and some cells may have endoreduplicated the remaining copy, leading to a mixed population with different genotypes. (C) By patching cells on a 5-FOA plate, the majority of cells containing the URA3 marker are killed, but a mixture of different cell types still potentially exists. (D) Tetrad dissection and spore genotyping after meiotic culturing are used to assess the population genotype as indicated (see Supplemental Table S1). i. Tetrads from monosomic cells should yield two Ura– spores and two dead spores lacking chrV. ii. Tetrads from cells that failed to destabilize the centromere and that escaped killing on 5-FOA should yield two Ura– and two Ura+ spores. iii. Chromosome missegregation in monosomic or endoreduplicated cells may yield tetrads with one or three Ura– spores, respectively. iv. Endoreduplication should yield tetrads with four Ura– spores.

**Supplemental Figure S2.**
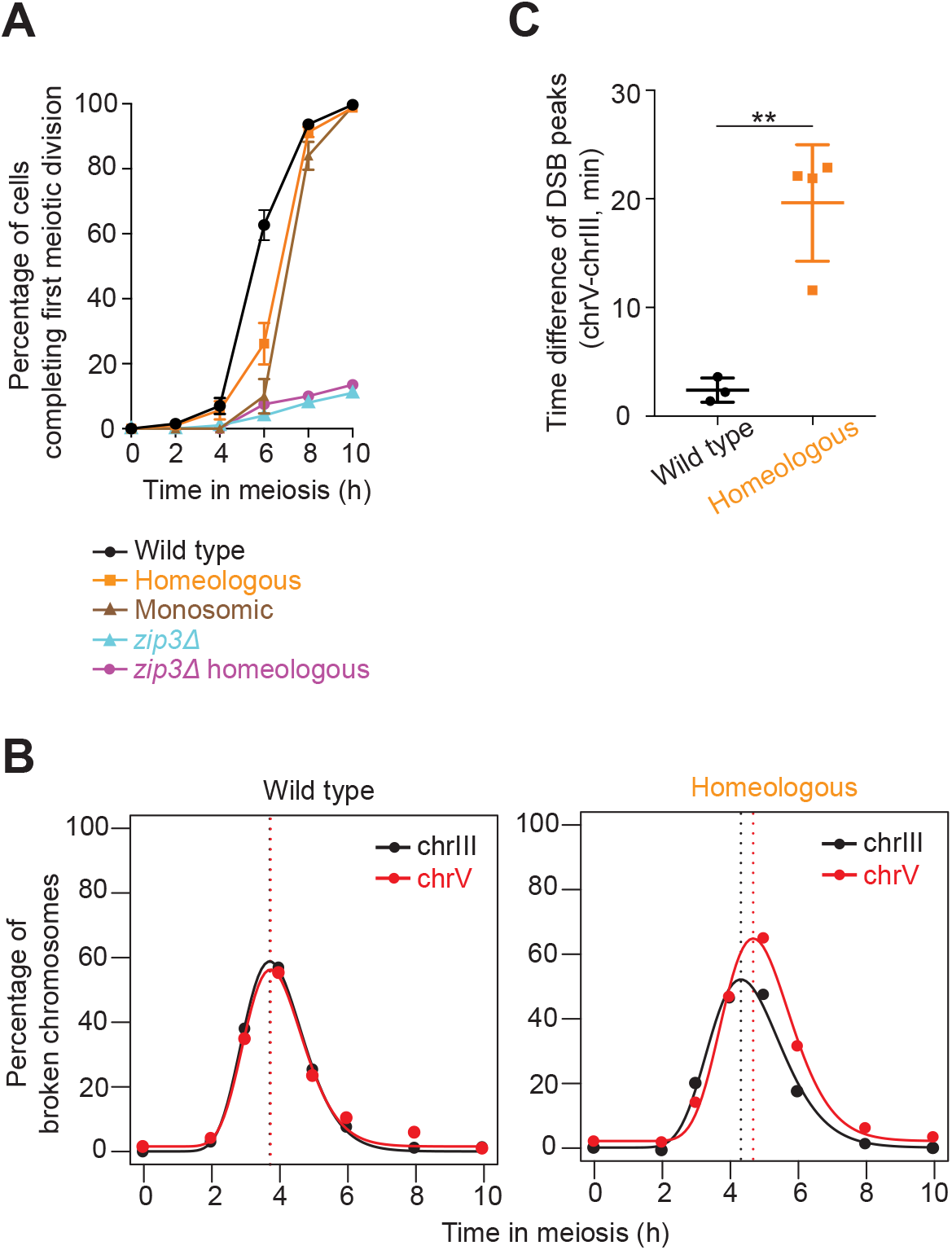
A later DSB peak on homolog-engagement defective compared to proficient chromosomes. (A) Meiotic progression showing percentage of cells completing the first division (total bi- and tetra-nucleate cells). ≥100 cells were counted at each time point for each sample. Error bars are mean ± SD for 3 wild type and 4 homeologous cultures; mean ± range for 2 monosomic, 2 *zip3Δ*, and 2 *zip3Δ* homeologous cultures. (B) Representative lognormal curve fitting exercises for chrIII and chrV in wild type and homeologous strains. (C) Quantification of DSB peak time differences after curve fitting between chrV and chrIII for wild type and homeologous strains. Each point is an independent culture; bars are mean ± SD. ** p<0.01, unpaired t test.

**Supplemental Figure S3.**
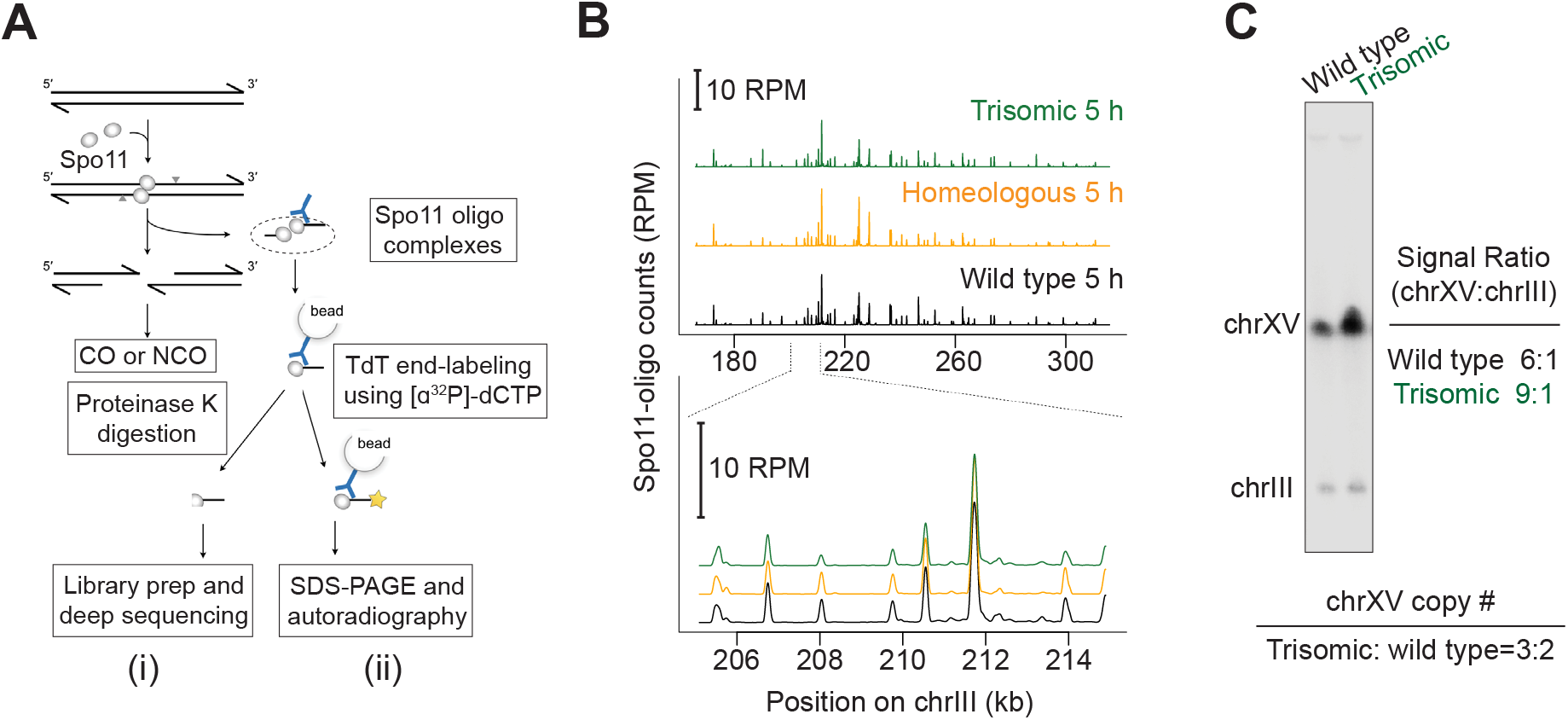
DSB mapping and quantification via analysis of Spo11-oligo complexes, and karyotypic analysis of the trisomy-XV strain. (A) Spo11 generates covalent protein-linked DSB; endonucleolytic cleavage releases Spo11 bound to short oligos. Spo11-oligo complexes can be detected by (I) Spo11-oligo mapping, in which immunoprecipitated Spo11-oligo complexes are digested with proteinase K and the released oligos are ligated to sequencing adaptors; or (II) labeling of Spo11-oligo complexes, in which immunoprecipitated Spo11-oligo complexes are radiolabeled using terminal deoxynucleotidyl transferase and [α-32P]-dCTP then separated by SDS-PAGE. (B) Spo11-oligo distributions on representative regions of chrIII. RPM, reads per million mapped; profiles were smoothed with a 201-bp Hann window. (C) Genotyping by Southern blotting to detect full-length chrXV and chrIII (control) separated by PFGE.

**Supplemental Figure S4.**
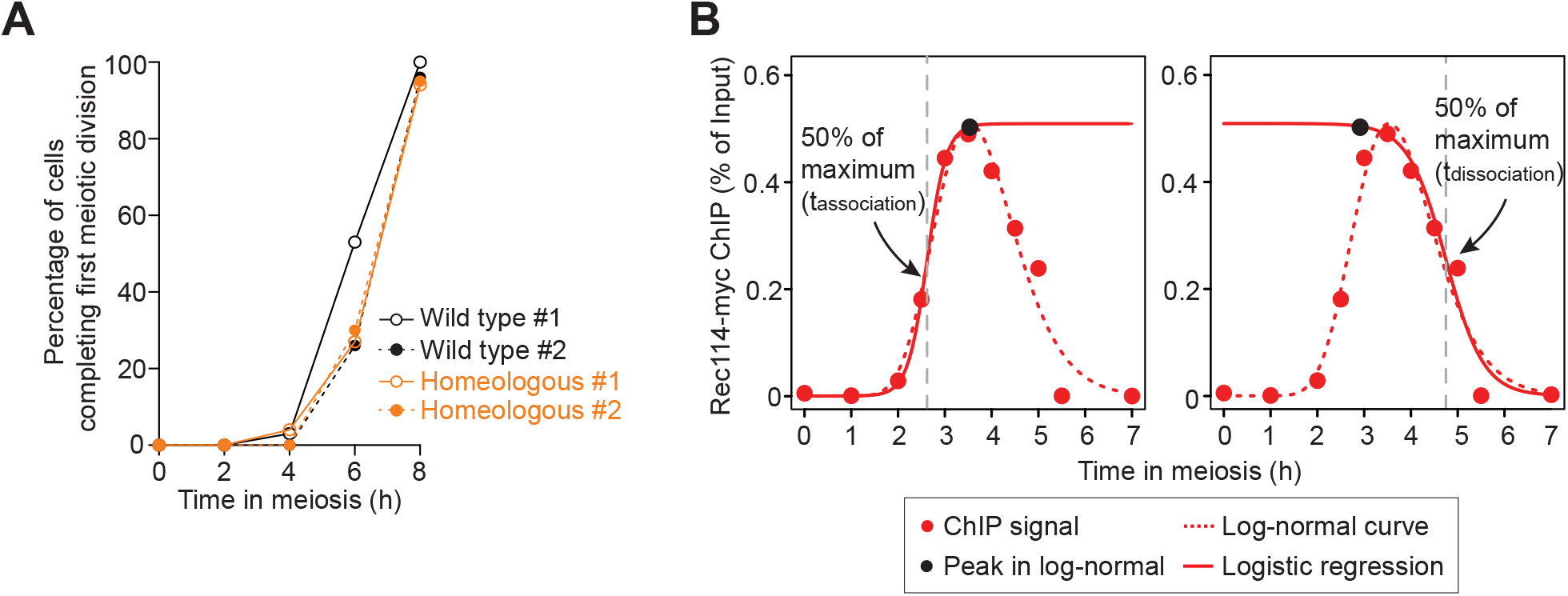
Time course analysis of Rec114 binding on homeologous chrV. (A) Meiotic progression showing percentage of cells completing the first division (total bi- and tetra-nucleate cells) for two independent data sets labeled as #1 and #2. At least 100 cells were counted at each time point for each sample. Note that wild type #2 appears slightly delayed. This exemplifies normal culture-to-culture variability in absolute timing of meiotic progression, highlighting the usefulness of internally controlled experiments. (B) Illustration of two-step curve fitting for calculations of association and dissociation times. A log-normal curve (dashed line) is first fitted to all points to define the peak time and height (black circle), then two logistic curves (solid lines) are fitted (left, using early points; right, using late points); t_association_ is defined as the position where the logistic curve reaches 50% of the peak level for early points (left) and t_dissociaiton_ is where the logistic curve reaches 50% of the peak level for late points (right).

**Supplemental Figure S5.**
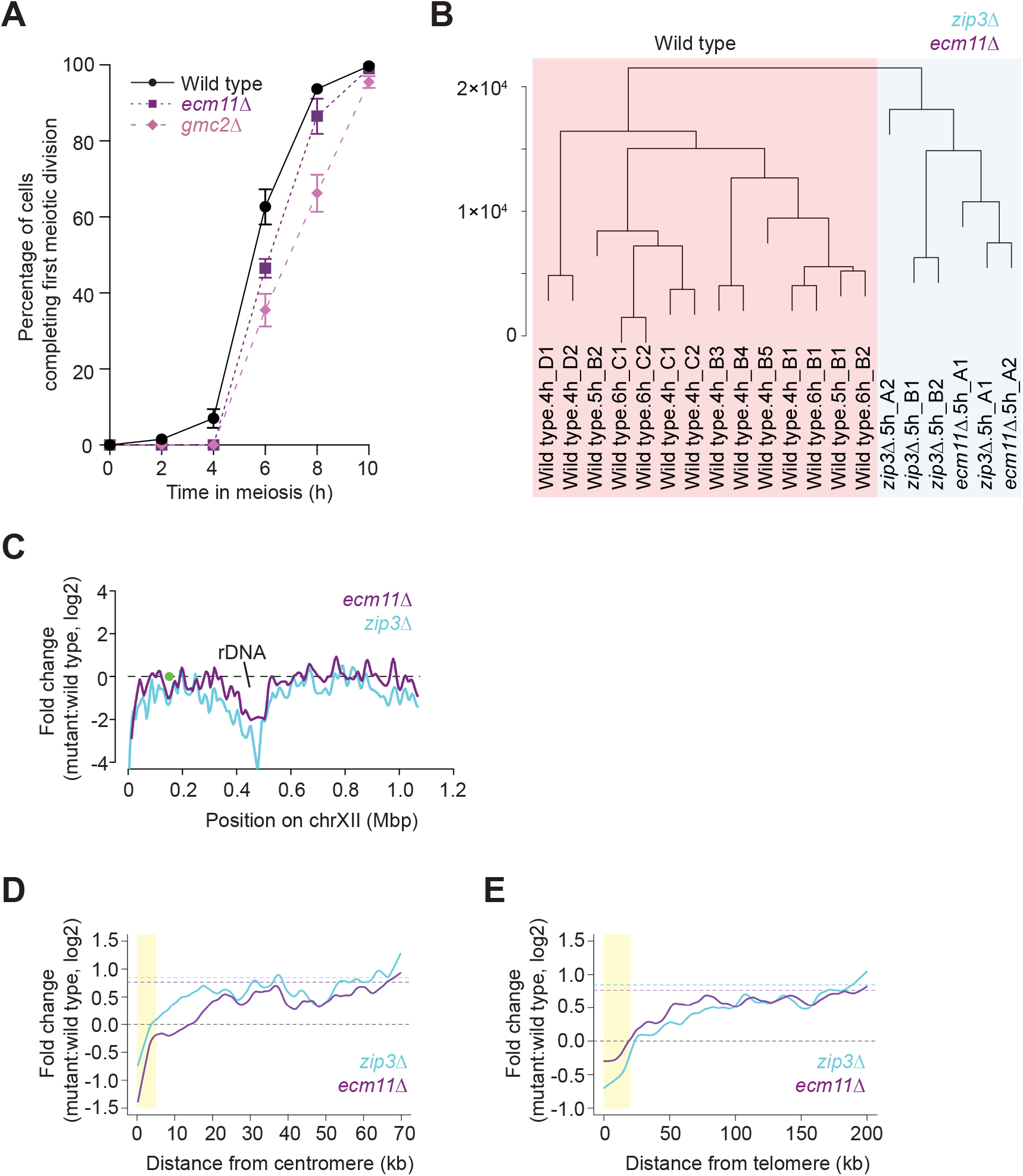
DSB patterns in gmc2Δ and ecm11Δ mutants. (A) Meiotic progression showing percentage of cells completing the first divisions (total bi- and tetra-nucleate cells, mean ± SD, 3 cultures each). At least 100 cells were counted at each time point for each culture. (B) Hierarchical clustering analysis using the “Ward D2” method for 21 wild-type and mutant maps. The y-axis represents height of the cluster dendrogram. In the label for each data set, the letters A, B, C and D stand for maps generated by Xiaojing Mu, Neeman Mohibullah, Megan van Overbeek, and Xuan Zhu, respectively, and the numerals distinguish biological replicate maps. (C) Regional variation in response to *zip3Δ* and *ecm11Δ* mutations for chromosome XII. Lines, local regression (loess) based on Spo11-oligo density changes in 5 kb-bins; green circle, centromere; rDNA location is labeled. (D, E) Fold change of Spo11-oligo densities for *zip3Δ* and *ecm11Δ* compared to wild type in pericentric (D) and telomere-proximal (E) regions. Lines, local regression (loess) for data in 5 kb-bins averaged across 32 chromosome arms; dashed line, no change in black and genome average change in cyan (1.8-fold, *zip3Δ*) and purple (1.7-fold, *ecm11Δ*); yellow shading, zones suppressed for DSB formation as previously defined (Pan *et al.* 2011).

**Supplemental Table S1.** Genotyping of monosomic biological replicates.

| Event | Observed phenotype | No. of viable spores | Culture <sup>a</sup> |  |
| --- | --- | --- | --- | --- |
|  |  |  | A | B |
| **Failure to kill *URA3* spores** | 2:2 Ura<sup>+</sup>:Ura<sup>-</sup> | 4 | 0 (0/13) | 0 (0/10) |
| **Endoreduplication** | Ura<sup>-</sup> | 4 | 0 (0/13) | 0 (0/10) |
| **Monosomic chrV** | Ura<sup>-</sup> | 2 | 84.6% (11/13) | 80% (8/10) |
| **Missegregation** | Ura<sup>-</sup> | 1 or 3 | 15.4% (2/13) | 20% (2/10) |

<sup>a</sup> The number of dissected tetrads is indicated:
Culture A: 13 tetrads dissected in total
Culture B: 10 tetrads dissected in total

**Supplemental Table S2.**
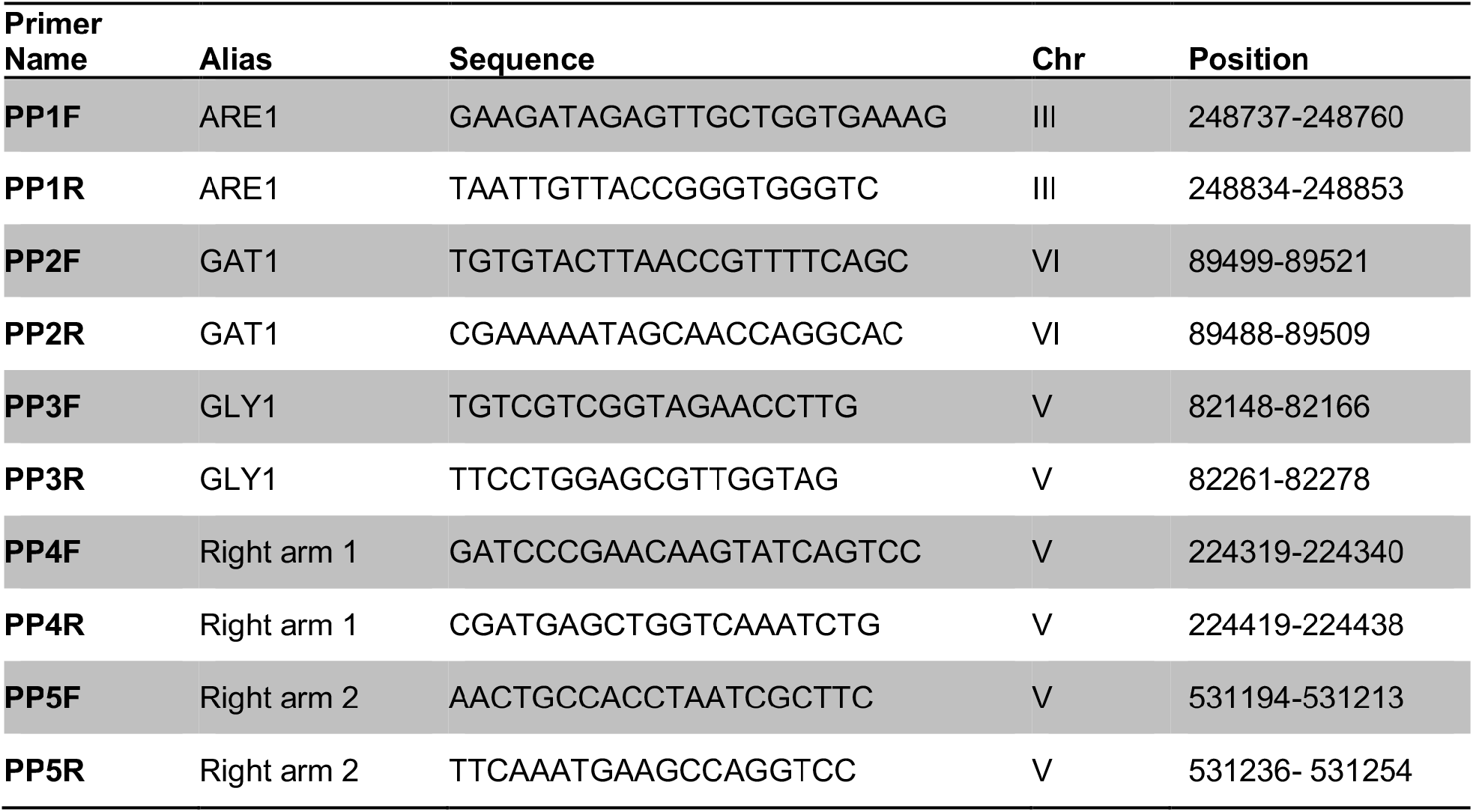
Primer pairs for ChlP-qPCR.

**Supplemental Table S3.**
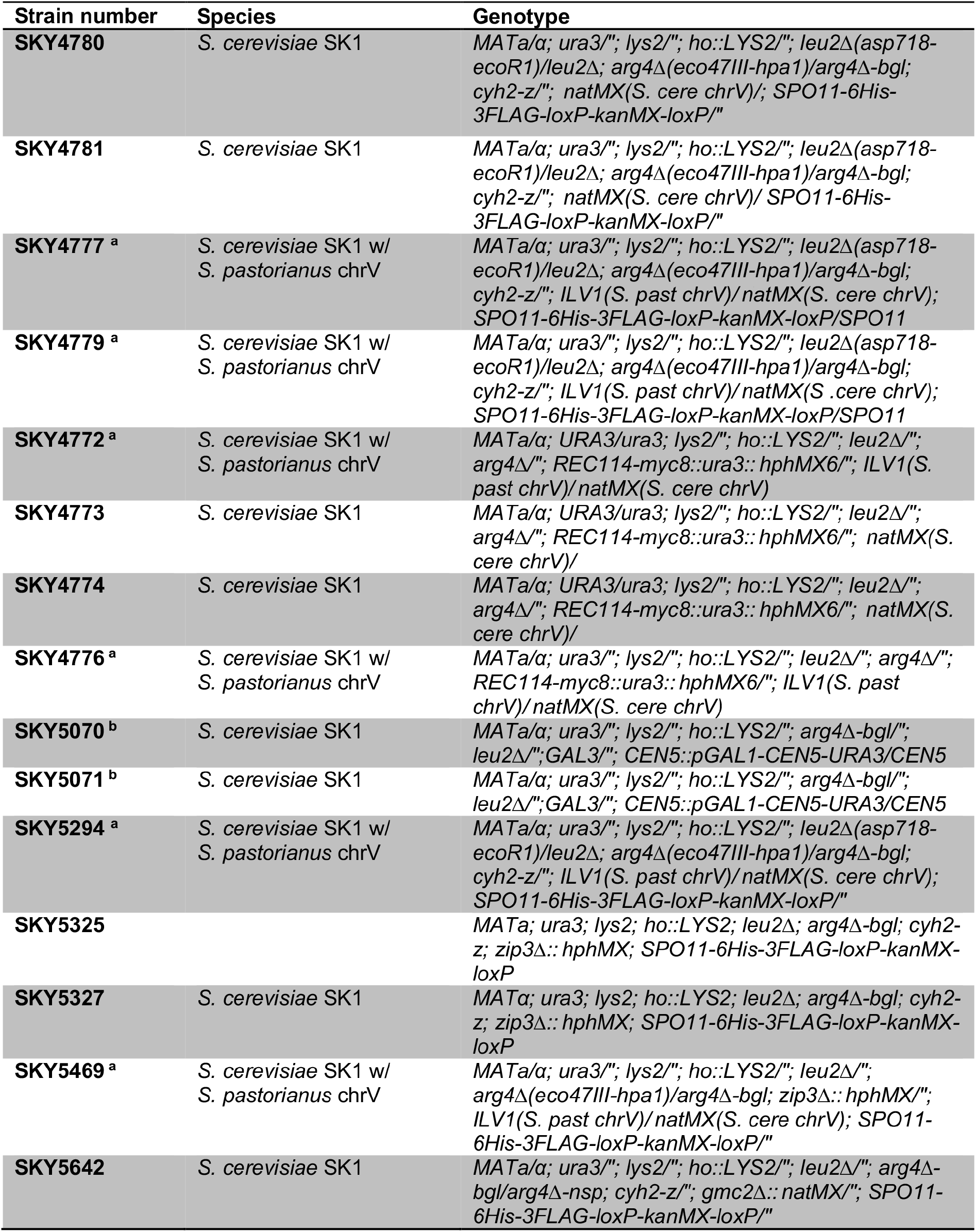
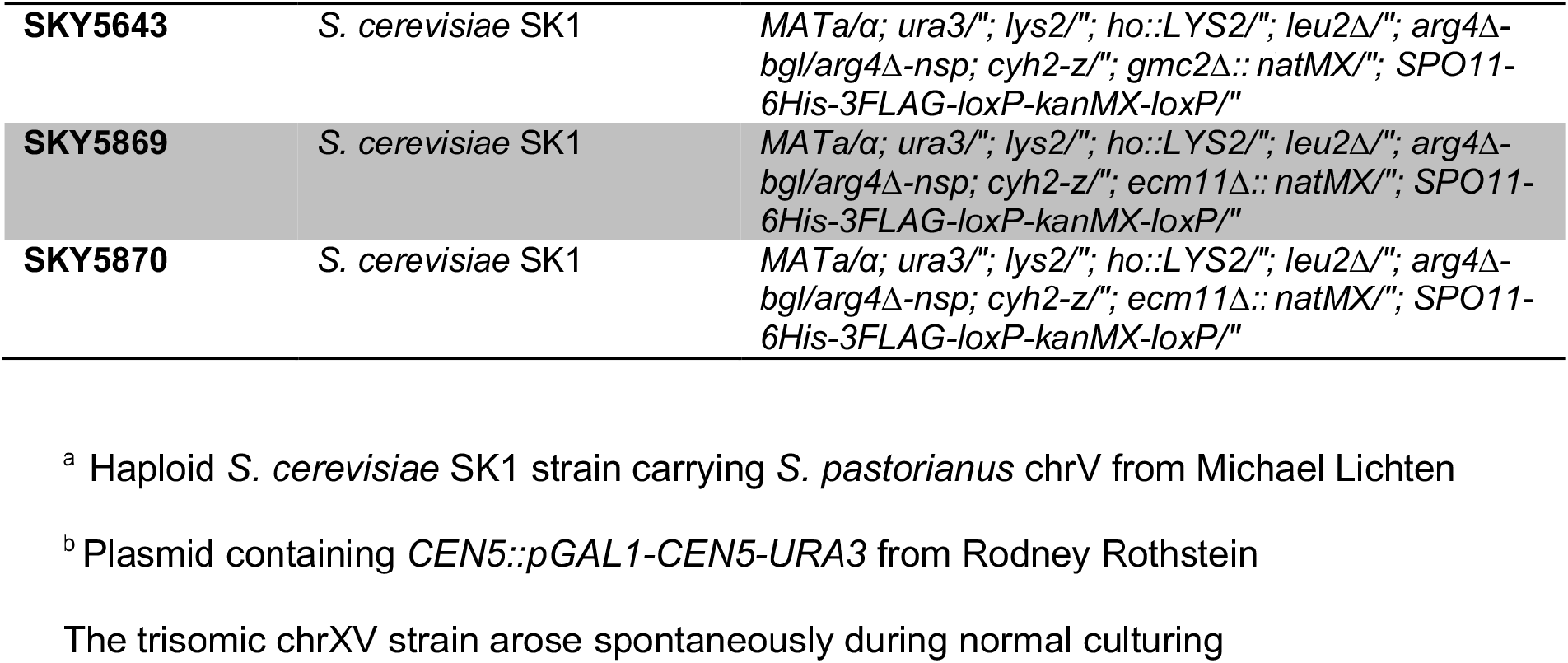
Yeast strains used in this study.

**Supplemental Table S4.** Mapping statistics for Spo11 oligo sequences.

| Dataset | Strain | Time | No. of total reads | Genome mapped to | No. mapped | No. mapped uniquely |
| --- | --- | --- | --- | --- | --- | --- |
| **Homeo_1<sup>a</sup>** | SKY5294 | 4 h | 19,996,390 | *S. cerevisiae* S288C plus *S. eubayanus* chromosome V | 17,815,499 | 15,000,784(98.1% of filtered) |
| **Homeo_2<sup>a</sup>** | SKY5294 | 5 h | 33,244,263 | *S. cerevisiae* S288C plus *S. eubayanus* chromosome V | 30,836,597 | 25,837,596(97.2% of filtered) |
| **zip3Δ_1<sup>b</sup>** | SKY5325x<br>SKY5327 | 5 h | 34,251,244 | *S. cerevisiae* S288C | 33,274,600 | 28,128,137(98.1% of filtered) |
| **zip3Δ_2<sup>b</sup>** | SKY5325x<br>SKY5327 | 5 h | 77,343,014 | *S. cerevisiae* S288C | 75,329,644 | 67,157,785(98.1% of filtered) |
| **zip3Δ homeo_1<sup>a</sup>** | SKY5469 | 5 h | 90,075,914 | *S. cerevisiae* S288C plus *S. eubayanus* chromosome V | 16,626,735 | 13,928,149(98.0% of filtered) |
| **zip3Δ homeo_2<sup>a</sup>** | SKY5469 | 5 h | 7,218,891 | *S. cerevisiae* S288C plus *S. eubayanus* chromosome V | 3,796,498 | 3,356,563(97.1% of filtered) |
| **ecm11Δ_1<sup>b,\*</sup>** | SKY5869 | 5 h | 10,778,802 | *S. cerevisiae* S288C | 10,648,118 | 1,435,809(98.0% of filtered) |
| **ecm11Δ_2<sup>b</sup>** | SKY5869 | 5 h | 9,679,255 | *S. cerevisiae* S288C | 8,086,230 | 7,254,361(98.0% of filtered) |

<sup>a</sup> One copy of chrV is from *S. pastorianus* and all other chromosomes are from the *S. cerevisiae* SK1 strain. Sequencing reads were mapped to the S288C genome (sacCer2 assembly) plus *S. eubayanus* chrV.
<sup>b</sup> Strains are *S. cerevisiae* SK1 background, mapped to the sacCer2 assembly.
\* For this sample, a synthetic oligonucleotide (TGGATCCGTGGAATAATGCACAATT) was added to increase the DNA amount during library preparation.

**Supplemental Table S5.** Spo11-oligo datasets used in this study.

| Source<br>(accession number) | Dataset<br>as shown at GEO | Alias used in<br>Supplemental<br>Figure S5 | Time | Tag | Figures |
| --- | --- | --- | --- | --- | --- |
| **Thacker et al. 2014**<br>(GSE48299) | WT_1 | Wild type.4h_B3 | 4 h | PrA | 3, 4 and 6; S5 |
|  | WT_2 | Wild type.4h_B4 | 4 h | PrA | 3, 4 and 6; S5 |
|  | zip3_1 | zip3Δ.5h_B1 | 5 h | PrA | 6; S5 |
|  | zip3_2 | zip3Δ.5h_B2 | 5 h | PrA | 6; S5 |
| **Mohibullah**<br>**and Keeney 2017**<br>(GSE84696) | wt4_1 | Wild type.4h_B1 | 4 h | PrA | 2 and 6; S5 |
|  | wt4_2 | Wild type.4h_B5 | 4 h | PrA | 6; S5 |
|  | wt5_1 | Wild type.5h_B1 | 5 h | PrA | 2 and 6; S3 and S5 |
|  | wt5_2 | Wild type.5h_B2 | 5 h | PrA | 6; S5 |
|  | wt6_1 | Wild type.6h_B1 | 6 h | PrA | 2 and 6; S5 |
|  | wt6_2 | Wild type.6h_B2 | 6 h | PrA | 6; S5 |
| **Murakami et al. 2020**<br>(GSE119689) | wt_a_4h | Wild type.4h_C1 | 4 h | flag | 6; S5 |
|  | wt_b_4h | Wild type.4h_C2 | 4 h | flag | 6; S5 |
|  | wt_a_6h | Wild type.6h_C1 | 6 h | flag | 6; S5 |
|  | wt_b_6h | Wild type.6h_C2 | 6 h | flag | 6; S5 |
| **Zhu and Keeney 2015**<br>(GSE67910) | WT_1 | Wild type.4h_D1 | 4 h | flag | 6; S5 |
|  | WT_2 | Wild type.4h_D2 | 4 h | flag | 6; S5 |
| **This study**<br>(GSE152957) | SK1_homeologous_4h |  | 4 h | flag | 2 and 6 |
|  | SK1_homeologous_5h |  | 5 h | flag | 2, 3 and 6; S3 |
|  | SK1_zip3Δ_1 | zip3Δ.5h_A1 | 5 h | flag | 4 and 6; S5 |
|  | SK1_zip3Δ_2 | zip3Δ.5h_A2 | 4 h | flag | 4 and 6; S5 |
|  | SK1_zip3Δ homeo_1 |  | 5 h | flag | 4 and 6 |
|  | SK1_zip3Δ homeo_2 |  | 5 h | flag | 4 and 6 |
|  | SK1_ecm11Δ_1 | ecm11Δ.5h_A1 | 5 h | flag | 6; S5 |
|  | SK1_ecm11Δ_2 | ecm11Δ.5h_A2 | 5 h | flag | 6; S5 |
|  | SK1_trisomic_4h |  | 4 h | PrA | 2 |
|  | SK1_trisomic_5h |  | 5 h | PrA | 2 and 3; S3 |
|  | SK1_trisomic_6h |  | 6 h | PrA | 2 |

